# Engineered retrovirus-like nanocarriers for messenger RNA delivery into neurons

**DOI:** 10.1101/2022.12.07.518870

**Authors:** Wenchao Gu, Sijin Luozhong, Simian Cai, Ketaki Londhe, Nadine Elkasri, Robert Hawkins, Zhefan Yuan, Kai Su-Greene, Margaret Cruz, Yu-Wei Chang, Patrick McMullen, Chunyan Wu, Changwoo Seo, Akash Guru, Wenting Gao, Tara Sarmiento, Chris Schaffer, Nozomi Nishimura, Richard Cerione, Melissa Warden, Robert Langer, Shaoyi Jiang

## Abstract

Systemic delivery of mRNAs into disease neurons is first limited by the blood-brain-barrier (BBB). Leukocyte-derived extracellular vesicles (EVs) can cross the BBB at inflammatory sites, emerging as promising carriers to target the disease brain. However, efficient mRNA loading into EVs and their uptake by neurons remain challenges. Here we incorporated inside EVs the endogenous retrovirus-like Arc protein capsids, stabilized by Arc 5’UTR RNA elements, to effectively load and deliver mRNAs. Produced from self-derived leukocytes, engineered retrotransposon Arc EVs (eraEVs) are immunologically inert with minimal clearance. Equipped with endothelial adhesion molecules from donor leukocytes, circulating eraEVs enter the brain enriching at neuro-inflammatory sites. During self-assembly, Arc recruits enveloping proteins onto eraEVs further promoting neuronal uptake. Possessing high effectiveness like viral vectors and biocompatibility as natural vesicles, eraEV-nanocarriers can be produced from virtually all donor cell types, potentially leading to the development of future clinical therapies for a range of diseases.

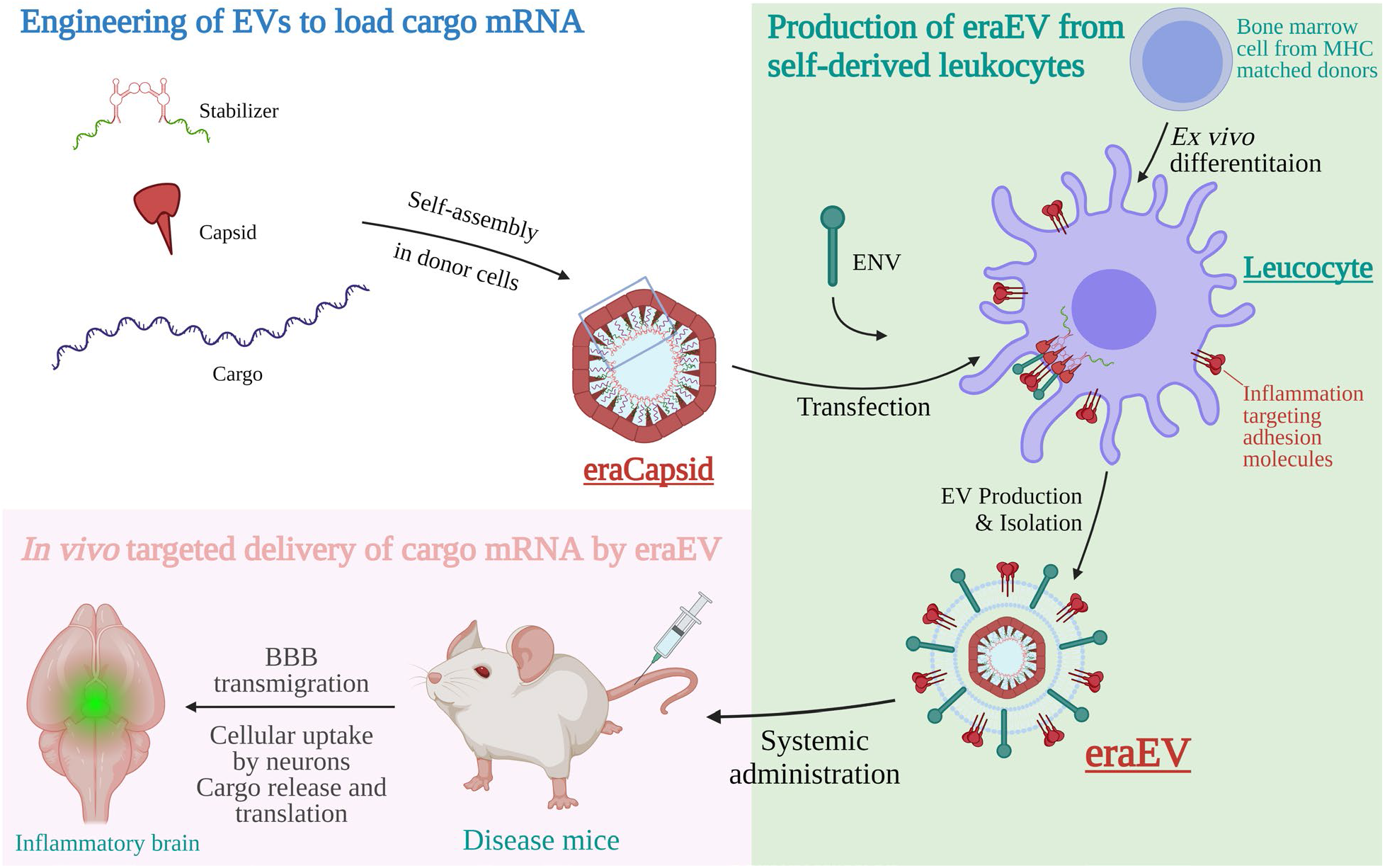

## Introduction

Systemic drug delivery into neurons is often limited by the BBB preventing drug entry into the brain, the low efficacy of carrier uptake by neurons and cargo release in neurons. Most biologic drugs, e.g., recombinant proteins, therapeutic antibodies, or nucleic acids, do not cross the BBB^1,2^. AAVs can deliver DNA into the neurons, whereas recombinant proteins and therapeutic antibodies can cross the BBB when conjugated to receptor mediated transporters (RMTs) of certain peptides, such as insulin or transferrin. Messenger RNA (mRNA) has emerged as a new class of therapeutic agents to prevent and treat various diseases. To function *in vivo*, mRNA requires safe, effective, and stable delivery systems that protect the mRNA from degradation, allowing cellular uptake of carriers, as well as intracellular release of cargoes. A variety of viral and non-viral carriers have been developed, including retroviral vectors, lipid nanoparticles, polymers, protein derivatives and cell-membrane enclosed vesicles. Still, effective and targeted *in vivo* delivery remains challenging for RNA-based therapies.

In response to inflammatory stimuli from the brain, the BBB becomes disrupted as brain microvascular endothelial cells exhibit increased permeability with elevated expression of leukocyte adhesion molecules^3-5^. These allow circulating leukocyte-derived EVs to enter the brain, making them great candidates as drug carriers to target neurons^6-10^. The BBB becomes more permeable to leukocyte EVs in various neurological conditions, including age-associated chronic inflammaging^11^, neuro-degenerative diseases^3^, and more severe pathological changes like systemic inflammation^4^ and secondary injury (e.g., stroke)^5^. However, the clinical translation of EV-based therapies for these conditions is hampered by the low cargo capsulation efficacy and a lack of control over which molecules are loaded into EVs from their donor cells. The cargo of EVs may include proteins, DNAs, RNAs, lipids, nutrients, and metabolic waste. Unwanted cellular components cannot be excluded from EVs, not only compromising the loading capacity, but also delivering potentially harmful components to the target.

Many systems were developed to load small RNAs, e.g., siRNAs and microRNAs into EVs, but active enrichment of long mRNAs in EVs remains a challenge^12^. Extremely low copy numbers of endogenous EV-associated RNAs were reported, ranging from 0.02 to 1 RNA per EV. Small RNAs are more efficiently packaged into EVs than long mRNAs (0.01 to 1 microRNA *vs*. 0.001 long intact RNA per EV)^13,14^. Only 8% of mRNAs in donor cells may be detected in their EVs^15^. Previous methods for loading mRNA into EVs include passive and active encapsulation^16,17^. For example, Catalase mRNAs were loaded by incubation with macrophage derived EVs after sonication and extrusion, or permeabilization with saponin to treat Parkinson’s disease^8^. Such post-EV-isolation loading methods have limited cargo capacity as EVs already full of donor cellular components cannot be unloaded to make space for drugs. EVs can also be engineered on the level of parental cells^16,18,19^. The ability to incorporate mRNAs efficiently and selectively into EVs under natural conditions is critical to the design of EVs as therapeutic drug carriers.

Widely distributed across the eukaryotic domain, transposable genomic elements of the long-terminal-repeat (LTR) retrotransposons and retroviruses have evolved from the same origin. A homolog of the capsid protein (Gag) of LTR retrotransposons and retroviruses, Arc (Activity-regulated cytoskeletal-associated protein) may have evolved in parallel and acts similarly to infectious RNA retroviruses, mediating intercellular communication in the central nervous system (CNS)^20^. PEG10, another Gag homolog, was recently pseudo-typed to form virus-like particles for mRNA delivery *in vitro*^21^. Arc proteins self-assembles into virus-like capsids to encapsulate mRNAs. The Arc capsid is released from neurons in EVs to mediate the transfer of mRNAs into recipient neurons via receptor mediated endocytosis^20^. Despite its great potential as a carrier, Arc EVs have been under-explored in drug delivery. Without their UTRs added, *Arc* and *Gag* were reported with low efficiency in mRNA transduction and little specificity for a particular cargo^22,23^. In our study, the addition of Arc 5’ UTR (A5U) to the cargo construct stabilized the capsid, increasing cargo loading capacity. Thus, we incorporated the Arc protein capsid into the lumen of EVs, together with the stabilizer A5U RNA motif, to effectively encapsulate and transfer mRNAs.

Produced from self-derived leukocytes, eraEVs are immunologically inert and actively enrich in inflammatory microenvironments transmigrating across the BBB with the help of donor leukocyte’s membrane molecules. Meanwhile, Arc components recruit enveloping proteins during self-assembly and subsequently promote cellular uptake of EVs by recipient neurons, based on their native functions in mediating inter-neuronal exchange of molecules. Moreover, the virus-like capsid makes eraEVs more stable than other engineered RNA-loading EVs, protecting the cargo from RNase degradation until its release is triggered. While taking advantages of these unique virus-like features, eraEVs are safe. They function as transient drug carriers without the ability to replicate, infect nor insert genetic information into the recipient genome^20^. Importantly, eraEVs can be produced from most donor cells to be applied to a wide range of biomedical applications, based on the native targeting ability of EVs from different cell types. This study is the first example of an endogenous virus-like system capable of loading and delivering mRNA *in vivo*, particularly into disease neurons via systemic administration.

## Results and discussion

### The RNA motif A5U stabilizes Arc EVs

To date, the Arc vesicle has yet to be developed into a drug carrier due to inefficient cargo loading and transfer. The Arc protein capsid naturally enriches mRNAs into EVs. In fact, the assembly of *Arc* capsids requires RNA^20^. It was suggested that intra-neuronal trafficking and inter-neuronal transfer of *Arc* mRNA are supported by *Arc* 3’UTR (A3U), which interacts with the nucleocapsid NC domain of *Drosophila* Arc^22,24,25^. However, mammalian Arc lacks the NC domain and the presence of two conserved introns and *cis*-acting regulatory elements in A3U contribute to its decay via the nonsense-mediated RNA decay (NMD) pathway^26,27^. Moreover, the dendritic targeting element (DTE) in A3U destabilizes mRNA independent of the NMD pathway. Therefore, the addition of the full-length A3U to the cargo construct may lead to transient cargo expression. On the other hand, Arc exhibits similar biochemical properties as its homologs, retroviral Gag proteins^20,28^, which encapsulate the long viral RNA genomes in the context of a large excess of cytosolic host RNAs^29^. Packaging of the 5’ UTR of the *HIV-1* genome depends upon Gag protein intact capsid CA domain lattice. Arc protein has a conserved CA domain that can interact with its 5’ UTR and may also bind to other RNAs nonspecifically through ionic interactions in its N-terminus. We therefore hypothesize that A5U can promote the efficacy of mRNA cargo loading. We later found that A5U mainly acts to stabilize the Arc capsid.

We used rat Arc (rArc) to produce Arc EVs, by transfecting a DNA vector encoding the Arc capsid and cargo mRNAs (w/ or w/o rat A5U) into human embryonic kidney HEK293 cells (Fig.1A). For simplicity, we label rA5U as the <u>stabilizer</u> and rArc as the <u>capsid</u> in all figures. Under optimized production conditions (Fig.S1-2 and data not shown), EVs were isolated from the supernatant culture medium and fluorescently labeled by antiArc-Alexa532 for an analysis of the Arc+ subpopulation via fluorescent nanoparticle tracking analysis (NTA), whereas total EVs were measured by light scattering from all particles (Fig.1B-C and Fig.S3A-E’). The Arc+ EVs appeared larger (or denser) than Arc‒ vesicles (Fig.1D and Fig.S3B, C’-E’). Interestingly, only Arc together with the A5U motif enabled a significant proportion of Arc+ EVs, showing a larger mean size of total EVs (Fig.1E-F and Fig.S3A-B). Since the measurements were taken after a long period of EV production, collection followed by antibody staining, the low copy number of Arc+/A5U‒ EVs may be a result of decreased capsid stability rather than a low assembly rate. The 5’ UTR of *Arc* may function similarly to that of HIV, whose dimerization initiations sites (DIS) can dimerize and subsequently stabilize with the Gag protein nucleocapsid (NC) and capsid (CA) domains, packaging into assembling virions ^30.31^. Consostent with fNTA, epifluorescence reading of Arc EVs (antiArc-Alexa532+) and total EVs, labelled by a plasma membrane stain, CellMask Deep Red (CMDR), confirmed the relative quantification of EV subpopulations (Fig.S3F). Altogether, these suggest that A5U is required for the assembly of stable Arc Evs.

**Fig. 1.**
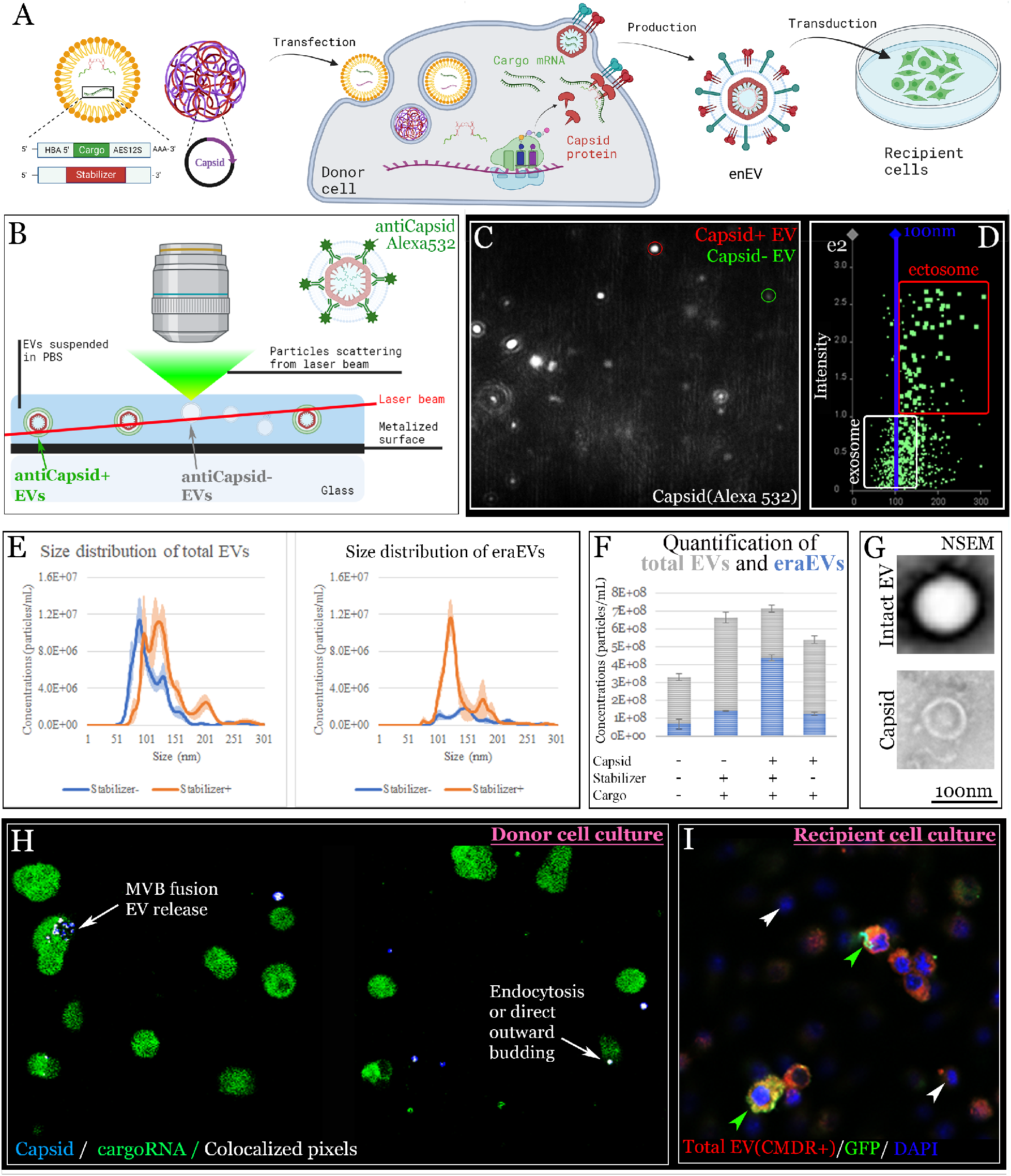
EV analyses indicate the importance of A5U for the assembly of stable Arc EVs. **(A)** The production of eraEV from donor cells and its transduction into recipient cells. **(B)** Fluorescent NTA measures the concentration and size distribution of antiArc-Alexa488+ EVs among all light-scattering EVs (screenshot in **C**: red, Arc-EV; green, non-Arc-EV). **(D)** Sizes of particles are plotted as a function of their scattered intensity. Arc-EVs appear larger than non-Arc-EVs. **(E)** Size distribution is represented as a histogram of particle concentrations of the EVs w/ or w/o the stabilizer. A lower camera level was used to measure fluorescent Arc+ EVs, with total EVs measured by a higher camera level. **(F)** Concentrations of total (gray) and Arc+ EVs (blue) are plotted. **(G)** NSEM of an Arc EV and a bare Arc capsid. **(H)** Confocal microscopy captured exosome release via MVB membrane fusion, and direct outward budding and/or endocytosis of EVs in donor cells. Arc protein expression is weak intra-cellularly, whereas bright extracellular vesicles were often seen, likely assembled Arc capsids. **(I)** Overlap between GFP and CMDR was observed in recipient cells: DAPI+/CMDR-cells w/o EVs showed no GFP signal.

We further characterized the morphology of Arc EVs via negative stain electron microscopy (NSEM), capturing both enveloped EVs and bare Arc capsid (Fig.1G). To visualize the secretion and transfer of Arc EVs, the Arc protein and cargo mRNA were labelled by immunocytochemistry and fluorescence *in situ* hybridization with quantitative hybridization chain reaction, followed by confocal microscopy (Fig.1H). In addition to many extracellular Arc vesicles, direct outward budding (also possibly endocytosis) of capsid+/stabilizer+ EVs was frequently observed, whereas the release from multivesicular bodies (MVBs) of the capsid+/stabilizer‒ EV subpopulation was more often seen. EVs have been classified into ectosomes and exosomes based on their size and biogenesis differences. Like other Gag proteins, Arc may mediate direct outward budding of ectosomes, which are larger than exosomes that are released from the MVBs fusing to donor cell membrane. This agrees with NTA size distribution, suggesting that capsid+/stabilizer+ Arc EVs are mainly ectosomes (Fig.1D-E and Fig.S3B, C’-E’). Moreover, engineered EVs successfully transferred cargo mRNAs to recipient cells *in vitro* (Fig.1I). The cargo was only translated in recipient cells that endocytosed CMDR+ EVs, but not all CMDR+ recipient cells expressed the cargo. The transduction efficacy highly depends on the donor-recipient cell pairs. Here we produced EVs from donor human HEK293T and transferred them to recipient human triple negative breast cancer cells (MDA-MB-231) or human glioblastoma cells (U-87 MG). Thus, we engineered, produced, and isolated Arc EVs that can load and transfer mRNA *in vitro*.

### Arc and A5U promote mRNA loading into EVs

We next investigated whether adding the capsid and stabilizer could improve the efficiency of cargo loading. We first fluorescently labelled mRNA transcripts to visualize cargo-loaded EVs in donor cells via live-cell imaging. From cargo constructs w/ or w/o A5U, *A5U*/*GFP* or *GFP* mRNAs were transcribed *in vitro*, using fluorescent Cy3-UTP or Cy5-UTP respectively to replace 25% of total UTPs (RNA QC analysis in Fig. S4). Co-transfecting combinations of these mRNAs with Arc DNA plasmids, we produced EVs of six control and experimental groups: (1) mock transfection (NT/NoTrans); (2) Capsid; (3) _*cargo*_*GFP*; (4) *stabilizer-*_*cargo*_*GFP*; (5) Capsid/_*cargo*_*GFP*; (6) Capsid/*stabilizer-*_*cargo*_*GFP*. To quantify and compare cargo loading into total EVs, we took Cy3/Cy5 fluorescence intensity reading of purified EVs, showing that Arc significantly elevated cargo loading by ∼6 fold (Fig.2A&D). Consistently, epifluorescence microscopy of donor cells showed a drastic increase in the amount of vesicle-size Cy3+/Cy5+ mRNAs (Fig.2B-C & 2E-F), likely encapsulated in Arc+ EVs as the Arc‒ control showed sparse extracellular signals. Previously, we could not determine whether A5U is required for the formation or preservation of Arc capsids, as analyses in Fig.1 were performed after a long period of EV processing, during which instable EVs may disassemble. Now, real-time live-cell imaging of donor cells shows that Arc can form capsids encapsulating cargo w/o A5U (Fig.2D&F).Altogether these suggest that <u>A5U is required for the stability of Arc EVs</u> rather than assembly. Moreover, <u>Arc significantly increases mRNA loading into the EVs</u>.

**Fig. 2.**
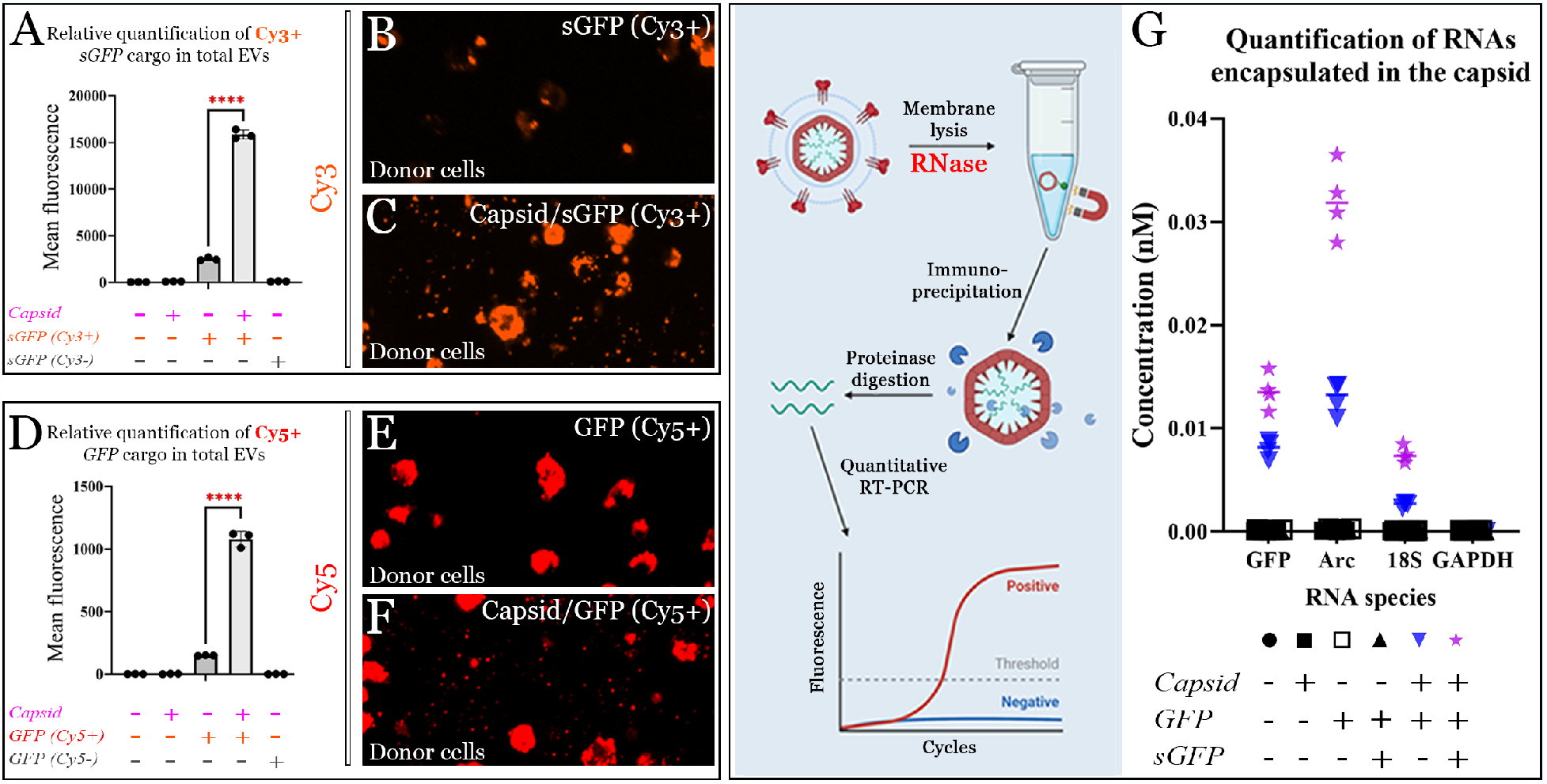
A5U increases mRNA encapsulation into EVs and Arc capsids in donor cells. **(A-F)** sGFP = A5U-GFP (A5U upstream of GFP in the same construct). The amount of Cy3+ *A5U-GFP* or Cy5+ *GFP* in EVs was increased in the presence of Arc capsid, shown by fluorescence intensity reading of purified EVs (**A&D**, quantification based on n=3 with ****P < 0.0001), as well as real-time epifluorescence imaging of the donor cells **(B-C&E-F). (G)** RIP followed by RT-qPCR showed that the A5U stabilizer increased mRNA loading into Arc capsids. Quantification based on n=4 with **P < 0.01.

We further investigated how A5U impacts mRNA encapsulation into total EVs and Arc capsids. The incorporation rates of different cyanine dyes vary in each mRNA transcript. Thus, to fairly compare the loading of Cy3-A5U+ *vs*. Cy5-A5U‒ cargos, we first measured fluorescence intensity of the two mRNAs at serial dilutions with known concentrations to create linear standard curves for absolute quantification (Table S1). We then transfected both mRNAs into the same donor cell culture and found that the amount of Cy3-A5U+ loaded into EVs was 1.48 folds of Cy5-A5U‒. This ratio was similar when two mRNAs were transfected separately (1.31:1) at the beginning of EV production, but the amount of A5U‒ cargo in the donor cell culture and in collected EVs declined rapidly over time. We next investigated mRNA encapsulation into Arc capsids, performing RNA immuno-precipitation (RIP) followed by RT-qPCR (Fig.2G). EVs were lysed followed by anti-Arc immuno-precipitation and RNase/DNase treatment to remove nucleic acids outside the capsids. Precipitated Arc capsid was then digested to release its cargo for absolute quantification by RT-qPCR using the standard curve method. The result greatly varies when the ratio between the capsid and cargo changes. At the molar ratio Capsid:*Cargo*=3:2, the presence of A5U increased the encapsulation of various RNAs into the capsid but not selectively of the A5U-GFP (Fig.2G). An abundant level of the capsid proteins is important for sufficient cargo loading (Fig.S5C-D). Thus, A5U <u>does not significantly contribute to the selectivity of cargo loading.</u>

### A5U+ Arc EVs improve the delivery efficacy and stability of cargo mRNAs

With fluorescently labelled mRNAs, <u>the efficacy and stability</u> of A5U+ Arc EVs as carriers were verified by drastic and stable increase of cargo mRNA transfer into recipient cells (Fig.3A). W/o A5U, cargo transfer was less abundant with lower stability, with a mild elevation of cargo accumulation for 3 days (1.5-fold peak on day 1) (Fig.3B), whereas the A5U+ group showed a 12-fold increase at day 6 (16-fold peak on day 3), both compared to the Arc‒ control (Fig.3A). This agrees with previous observation that Arc EVs may be less stable w/o A5U (Fig.1E-G). The mechanism may resemble HIV capsid assembly, which requires its 5’ UTR to stabilize Gag protein capsids^32^. Live-cell imaging showed that EVs carrying fluorescent cargo started docking on the recipient cell membrane as early as 15-mins after EV transfer (Fig.3A1-A2&B1-B2). From 1-hr, intracellular fluorescence was seen and by 4-hrs virtually all recipient cells received A5U+ Arc EVs (Fig.3A3-A4) while the uptake of A5U‒ Arc EVs was significantly less efficient (Fig.3B3-B4). The relative quantification was normalized by the 4-hr mock transfection group background levels of Cy3 or Cy5 fluorescence. The levels of stabilizer+ cargo mRNAs remained higher in the recipient cell culture over one week, with not only had more cells the cargo but also more cargo in each cell. The same amount of total EVs were loaded and entered recipient cells at about the same rate. To quantify their uptake rates, we used unlabeled mRNAs as cargos and CMDR to label total EVs, for their visualization and quantification. With an optimized dye concentration (Fig.3C-middle), serial dilutions of EVs with known concentrations can be accurately quantified in this assay (Fig.3D & S6). We counted a similar percentage of CMDR+ recipient cells among all sample groups at various time points after EV transduction (Fig.3E). This confirms that the improved *in vitro* mRNA delivery efficacy by the capsid and stabilizer is not just from an increase in total EVs added to the recipient cells. To conclude, stabilizer+ Arc EVs significantly improved the efficiency and stability of mRNA delivery *in vitro*.

**Fig. 3.**
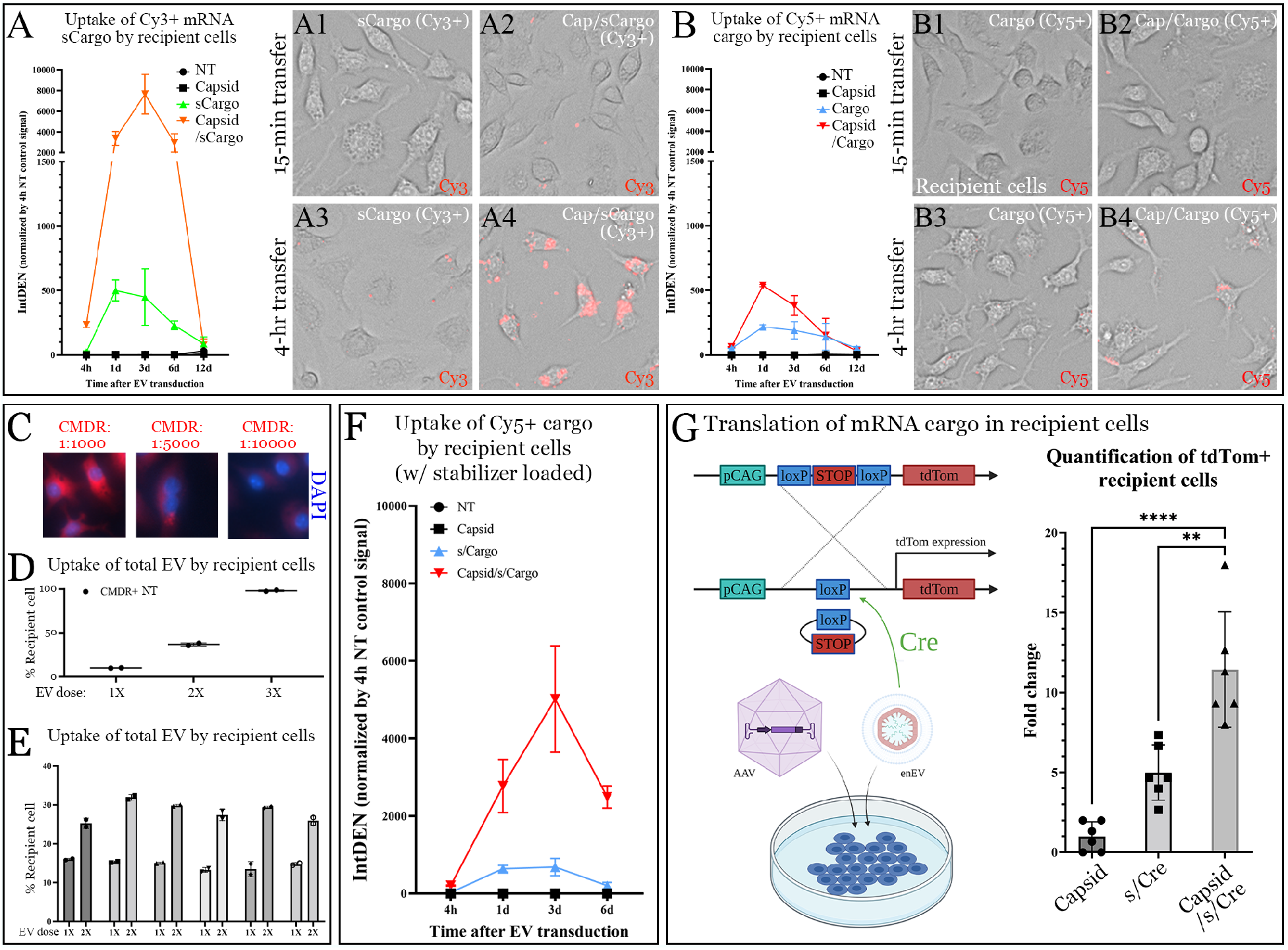
A5U enables effective and stable mRNA accumulation and translation in recipient cells. *sCargo:* A5U-*Cargo* (A5U upstream of *Cargo* sequence in the same construct). *s/Cargo*: A5U and *Cargo* both transfected but separately in two different transcripts. **(A)** Upon transfer of HEK293T-derived Arc EVs w/ (Cy3+) or w/o (Cy5+) the stabilizer, live-cell fluorescence imaging was performed at 15-min **(A1-A2)**, 4-hr **(A3-A4)**, 3-day, 6-day and 12-day time points to quantify mRNA uptake by RAW264.7 recipient cells. Integrated density of Cy3/Cy5 was normalized by background levels in 4-hr NT controls. The transfer of capsid+/stabilizer+ EVs showed a stable increase in cargo accumulation **(A**, orange), whereas capsid+/stabilizer‒ EVs delivered less efficiently with low stability **(B**, red**). (C)** CMDR was used to label total EVs. Titration experiments optimized the dye concentration to 1:5000, for moderate fluorescence with endosomal EVs visible in recipient cells, and near-linear correlation between cellular CMDR levels and doses of total EVs added **(D)**. As 3XEVs (3 donor cells *vs* 1 recipient cell) led to near-saturated signals (98.9% CMDR+ cells), we transferred 1X and 2X EVs to quantify total EVs endocytosed, observing a similar percentage of CMDR+ cells among all sample groups at 2 hours after EV transduction **(E). (F)** When the Cy5+ stabilizer‒ cargo mRNA was co-transfected with the stabilizer, the Cy5 signal persisted for a longer period in the recipient cell culture after EV transfer. **(G)** AAVs expressing loxp-STOP-loxp-tdTom were first transduced into the MB-MDA-231 recipient cell culture, followed by the EV transfer. 7-days after the transduction of AAVs with EVs, we observed a significant increase in tdTom expression using eraEV/_Cargo_Cre, whereas the stabilizer+ _cargo_Cre+ control also showed an in increase compared to other negative controls. Quantification of tdTom+ cells based on n=6. ****P<0.0001 and **P<0.01.

Despite a great advantage in mRNA transfer, we only observed a transient expression of cargo when the cargo mRNA construct contains an upstream A5U leading sequencing. Since our results suggest that the function of A5U is mainly to stabilize the capsid, we thus tested whether separating the A5U stabilizer from the cargo mRNA will enable and increase cargo expression. Indeed, when loaded together with an A5U stabilizer (w/ a reporter tail), the level of Cy5+ cargo mRNAs remained stably increased, and more importantly, the half-life of cargo expression was increased (Fig.3F). Thus, in following experiments, we co-transfect _cargo_mRNAs with the A5U stabilizer, with a reporter tail to monitor the transfection efficacy in donor cells. In fact, we found that the tail improves capsid stability, as A5U alone did not increase the proportion of Arc EVs among total EVs measured by NTA. This effect is unlikely to be sequence specific. If A5Us dimerize similarly as the HIV 5’ UTRs^30^, any downstream RNA sequence may non-specifically bind to the random coil structure on the C terminal of the Arc protein via ionic interactions and could further improve the stability of Arc capsids. As to the _cargo_RNAs, we added the 5’UTR of α-globin and 3’UTR of AES/12S for highly efficient protein translation. Such engineered EVs are termed as **eraEVs**, engineered retrotransponson Arc EVs. For better readout signals, we applied the high-throughput Cre reporting system^33^: (1) AAV transduction into recipient cells to express the loxP-flanked STOP cassette followed by a downstream tdTomato reporter; (2) the usage of Cre mRNA as the cargo (Fig.3G). We observed more tdTom+ recipient cells with eraEV/_cargo_Cre than the Arc‒ control (Fig. 3G). Therefore, eraEV can deliver mRNAs *in vitro* to be translated, with improved efficiency and stability.

### Leukocyte-eraEVs efficiently deliver mRNAs *in vivo* targeting neuro-inflammation

We next tested potential applications of eraEVs for neuronal diseases, as there are limited mRNA delivery methods into neurons^2^. BBB is a highly selective semi-permeable border to separate peripheral circulation from the CNS, composed of continuous brain microvascular endothelial cells (BMEC), their tight junctions, basement membranes, pericytes, and astrocyte terminals. BMECs normally express low levels of leukocyte adhesion molecules compared to peripheral endothelial cells to prevent the margination and transmigration of immune cells into the brain^34^. In healthy brains, circulating leukocytes move passively in the bloodstream along with the laminar flow of blood. At inflammatory sites, the hemodynamics of BBB becomes disrupted, greatly reducing the blood flow rate and increasing the chance that leukocytes adhere to BMECs, which now exhibit increased permeability and elevated expression of leukocyte adhesion molecules. These allow leukocyte-derived EVs to enrich in neuro-inflammatory micro-environments^6-10,35-37^. We first used the aging (also termed inflamm-aging) mice, as a low-grade chronic pan-neuronal neuro-inflammation model to test whether leukocyte-derived EVs can effectively enter the brain. We tested non-engineered EVs produced from a wide range of donor cells in the aging model (>90-week) and found that EVs from the primary bone marrow (BM) derived macrophages (MΦs) and dendritic cells (DCs) are the most potent ones accumulating in the brain, whereas EVs from immortalized cell lines and even other leukocytes are less efficient (Fig.4A). The BM-DC/MΦ EVs also enter the healthy steady-state brains, but mainly around the circumventricular organs.

**Fig. 4.**
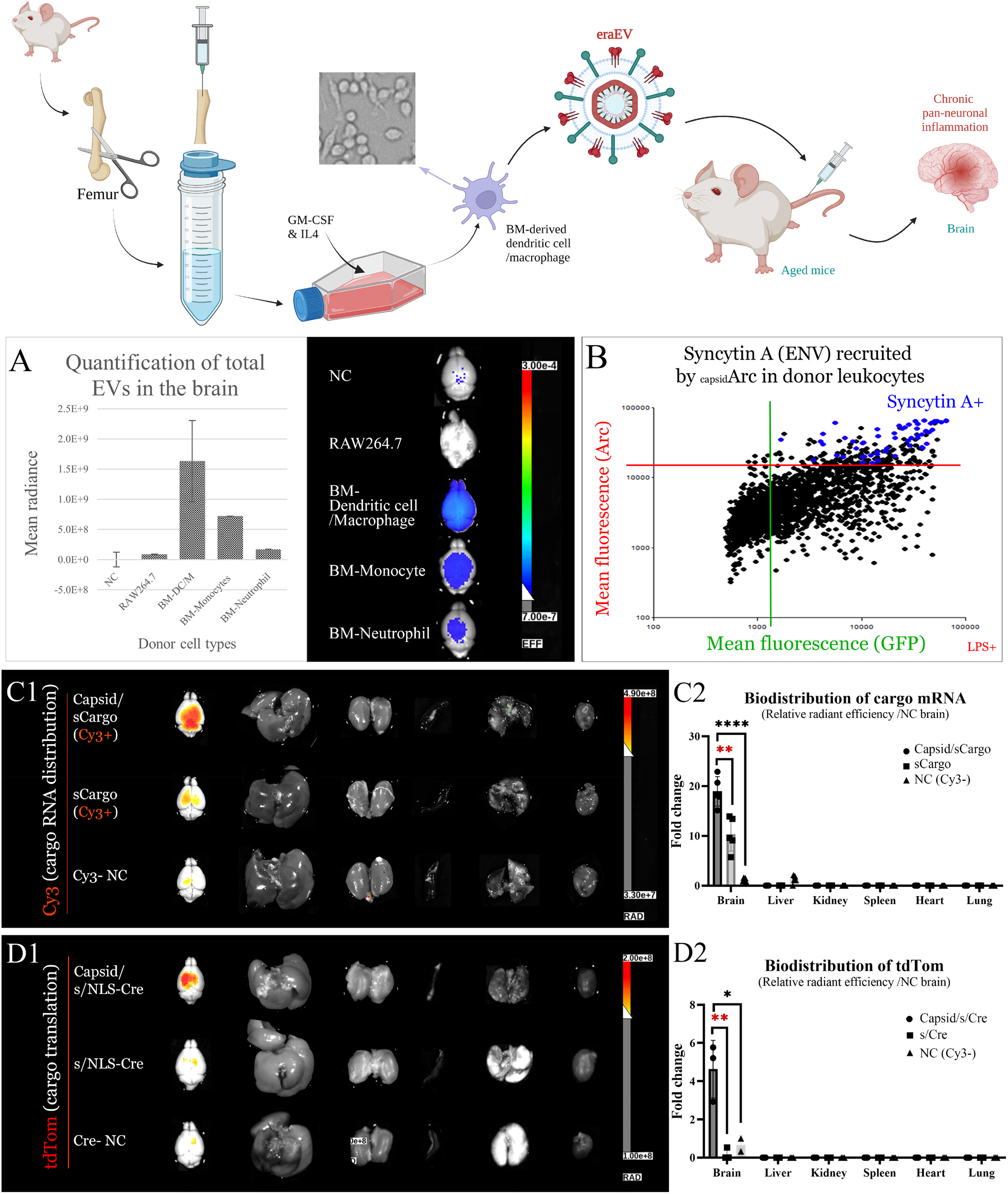
eraEVs deliver cargo mRNAs to be translated specifically into inflamm-aged brains. Among a variety of immortalized and primary donor cells tested, EVs produced from the bone marrow derived dendritic cells and macrophages showed the highest accumulation in the brain. An endogenous retroviral enveloping protein, Syncytin A, became upregulated by Arc overexpression in donor cells. Blue dots represent all Syncytin A+ cells in the donor cell culture. This population largely overlap with the Capsid+/Cargo+ donor cell population. **(C)** 3-days after IV injection of eraEVs: **(C1)** Arc+/Cy3-A5U+ EVs significantly enriched cargo mRNAs in aged brains, compared to Arc‒ and Cy3‒ controls. (Photo overlay of radiance displayed: color ranges from 3.3e+7 to 4.9e+8 and color threshold at 3.3e+8) **(C2)** Fold change of radiant efficiency: Mean±SD, n=5 in A5U/cargo or Arc/A5U/cargo groups; n=12 from various NC. **P<0.01 or **** P<0.0000001. **(D1)** Distribution of tdTomato indicates that mRNA cargo delivered into the brain was successfully translated from 3-days after IV injection. The expression was specific in the brain. (Color ranges from 1e+8 to. 2e+8 and a color threshold at 1.7e+8). Color thresholds were determined to subtract background signals based on negative control groups. **(D2)** Fold change of radiant efficiency: n=3 with *P<0.05 or **P<0.01. Mean±SD.

Leukocytes EVs can enter the neuro-inflammatory microenvironments, but without being uptaken by neurons they were rapidly metabolized and cleared from the brain. The native roles of Arc EVs in inter-neuronal mRNA transfer can further improve the neuronal uptake of these vesicles^20^, and eraCapsids can protect cargo mRNA from RNase degradation until the release is triggered. Interestingly, in donor leukocytes (monocytes and DC/MΦ), Syncytin A (SynA), an endogenous retroviral enveloping protein, appeared up-regulated by Arc overexpression especially when stimulated by LPS (lipopolysaccharide, Fig.4B). Thus, during the self-assembly of the Arc system, endogenous enveloping proteins may be recruited onto Arc EVs in leukocytes, and so we may not further introduce a cell fusogen component into our carrier system. Transfecting SynA did significantly promote the transduction rate in *in vitro* experiments involving certain donor-recipient cell pairs but did not improve *in vivo* delivery. It is possible that SynA overexpression interferes with transmigration across the BBB, a critical step during *in vivo* targeting.

We next tested leukocyte-eraEVs for *in vivo* delivery of mRNAs into the brain targeting the low-grade inflamm-aging. To produce a pool of immunologically inert EVs minimizing drug clearance *in vivo*, self-derived donor MΦs and DCs were differentiated *ex vivo* from BM cells extracted from mice of the same strain with a homogenous major histocompatibility complex haplotype (Fig.4&S7)^38^. As EVs from either MΦs or DCs were reported to cross BBB, we included all GM-CSF/IL4 differentiated BM culture subpopulations, including monocyte-derived DCs, monocyte-derived MΦs, and conventional DCs, most of which are in their progenitor state and LPS-activatable (Fig.S7A). For *in vivo* delivery, the same amount of total EVs among various sample groups was intravenously (IV) injected into inflamm-aged (>=90-week) mice. As described above, we add Cy3+ A5U-reporter RNAs to stabilize capsids and to monitor EV production. 3 days after the administration of control and experimental EVs, organs were extracted to be imaged via IVIS, following transcardial perfusion to remove signals in the circulating blood. We observed significant and specific enrichment of Cy3+ mRNAs in the brain via BM-DC/MΦ derived eraEVs (Fig.4C1-C2). Consistent with previous observation that Arc EVs may be less stable w/o A5U, Cy5 from the A5U‒/Arc+ EVs ended up mostly in the liver and kidney (Fig.S8). Thus, BM-DC/MΦ derived eraEVs effectively deliver mRNA into the brain targeting chronic, low-grade, pan-neuronal inflammation.

### BM-DC/MΦ-eraEV delivered mRNAs can be endocytosed by neurons and effectively translated

Finally, to verified protein translation from cargo mRNA, we produced BM-DC/MΦ eraEV/_*cargo*_*Cre* to deliver *NLS*-*Cre* into aged Ai14 mice, in which Cre recombinase, once translated from the cargo mRNA, can initiate robust tdTom expression (Fig.5A). IVIS of tdTom confirmed specific cargo expression in the brain, starting from 3-days after systemic administration (Fig.4D1-D2) and becoming more significant at 7-days (Fig.S9A). Thus, eraEVs specifically enrich cargo mRNA to be expressed in inflammaging brains *in vivo*. Still at 3-days after IV injection, the CMDR levels were measured via IVIS, showing comparable amounts of total EVs among sample groups. By this point, the dye accumulated mostly in the liver and kidneys, likely a result of systemic clearance (Fig.S10). We then examined neuronal accumulation of eraEVs and cargo expression at cellular resolution via confocal microscopy of cryo-sectioned brain slices. In the EV‒/CMDR+ control, no CMDR was seen in the brain (Fig.5B1-B3). The Arc+/cargo‒ control EVs and eraEVs showed comparable levels of CMDR on and beyond BMECs (Fig.5C-D&D4). But only eraEV/_cargo_Cre enabled robustly increased expression of tdTom (Fig.5D-F). Thus, BM-DC/MΦ eraEVs can deliver mRNA to be endocytosed by neurons, released and translated intra-neuronally.

**Fig. 5.**
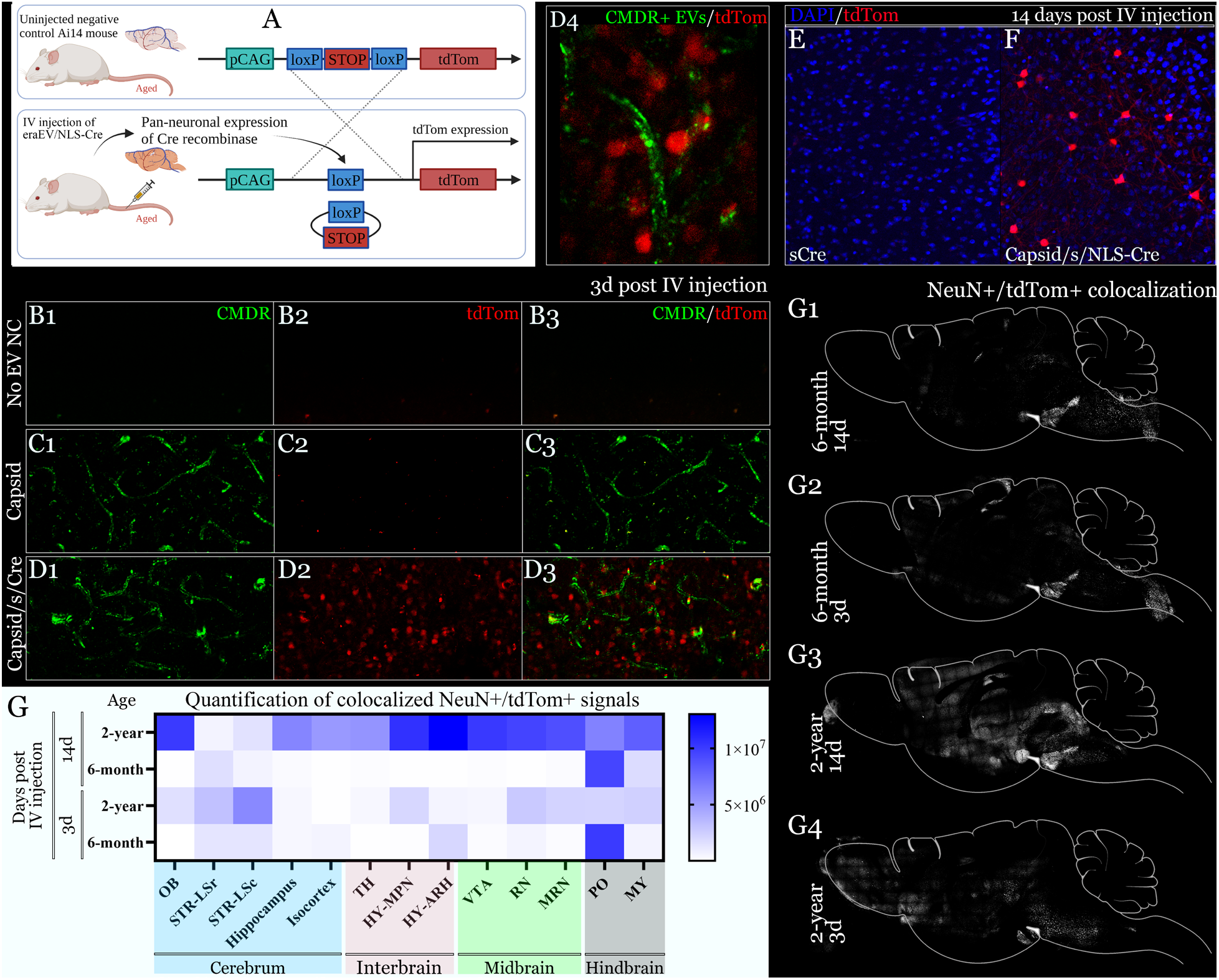
BM-DC/M eraEVs deliver mRNAs into neurons to be translated, across the BBB targeting chronic pan-neuronal inflammation. **(A)** BM-DC/M eraEVs/_cargo_Cre were intravenously injected into aged Ai14 mice. Arc+/A5U+/Cre+ EVs enabled tdTom expression in the inflammaging brain (D), distinct from the control groups (B-C). CMDR dye was seen on and beyond BMECs in both eraEV and Arc-control groups (C1, D1 and D4), but only eraEVs enabled tdTom expression (D, 3-days post administration; F, 14-days). **(G)** 14-days after eraEV/_cargo_*NLS-Cre* administration, aged brains showed a significant increase in NeuN+/tdTom+ expression in various brain regions (heat map plotted based on integrated density). OB, olfactory bulb; LSr, lateral septal nucleus, rostral; LSc, lateral septal nucleus, caudal; TH, thalamus; HY, hypothalamus; MPN, medial preoptic nucleus; ARH, arcuate hypothalamic nucleus; VTA, ventral tegmental area, RN, red nucleus; MRN, midbrain reticular nucleus; PO, pons; MY, medulla.

Next, to study the specificity of neuronal uptake of eraEVs, they were retro-orbitally administered to deliver *NLS-Cre* into aged and young Ai14 mice, whose brains were collected, fixed, thin-sectioned and imaged 3 or 14 days after the administration. Neurons were labeled by a neuronal marker, NeuN (Fox-3, Hexaribonucleotide Binding Protein-3), via immune-histochemistry staining. White pixels in Fig.5G1-G4 represent the colocalization between NeuN and tdTom, indicating neuronal expression of Cre. A higher level of neuronal tdTom was seen in aged brains compared to young controls in various brain regions, especially boundary regions surrounding circumventricular organs where the BBB is leakier even in the steady-state healthy brain (Fig.5G), whereas many tight-junction cerebrum and interbrain regions were also penetrated (e.g., striatum and isocortex). Altogether, taking advantage of the inflamm-aging model, we demonstrated *in vivo* <u>mRNA delivery by eraEVs</u> targeting the low-level chronic inflammation across the whole brain.

In conclusion, we engineered a safe and effective mRNA nanocarrier with *in vivo* targeted delivery. We found that an RNA regulatory motif, A5U, is required for the assembly of highly stable Arc EVs, which can encapsulate and transfer cargo mRNAs with higher efficacy. The incorporation of Arc itself already enables a dramatic increase in mRNA loading into total EVs, whereas the addition of A5U stabilizes the Arc capsids, further enriching cargo mRNA packaging. Combining Arc w/ A5U, we enabled highly abundant and stable accumulation of mRNA in recipient cells *in vitro*. Moreover, we demonstrated *in vivo* targeted delivery of mRNA via leukocyte-derived eraEVs, into the neurons of brains with chronic pan-neuronal inflammaging. Not only being the first case of an endogenous retroviral capsid used as a mRNA drug carrier with *in vivo* evaluations, but this is also the first example of effective *in vivo* targeted delivery of long mRNAs into neurons from the peripheral blood. This work highlights the potential of eraEVs to be used in novel therapies for inflammatory conditions in the CNS. The eraEV/cargo system can be produced from virtually all cell types, potentially leading to the development of future clinical therapies for a wide range of diseases.

## Methods

### Biological resources

An Arc plasmid (pGEX-6p1-GST-ArcFL, Addgene plasmid #119877) and a GFP plasmid (pcDNA3-EGFP, Addgene plasmid #13031) were purchased from Addgene. The rat Arc 5’ UTR sequence was synthesized by Genscript. Molecular cloning techniques (PCR amplification, restriction enzyme digestion and Gibson assembly) were used to generate constructs pCDNA3-Arc and pCDNA3-A5U-GFP. Correct sequences of all plasmids have been verified by Sanger Sequencing at Cornell Institute of Biotechnology or by Genewiz.

### *In vitro* transcription of DNA plasmids and RNA transcripts

Arc DNA plasmids were transfected into the donor cell culture via PEI or Xfect transfection reagent (Takara) following manufacturer’s instructions. DNA plasmids of pcDNA3-eGFP and pcDNA3-A5U-eGFP containing T7 promoters were linearized using PCR with a forward primer (CGC AAA TGG GCG GTA GGC GTG) upstream of the T7 (AG) promoter and a reverse primer (120*T TAG AAG GCA CAG TCG AGG) to add a 120 bp polyA sequence. The template of A5U stabilizer w/o a reporter tail was generated by PCR using a T7 forward primer, and a reverse primer w/o polyA added. These linearized templates were used to direct *in vitro* transcription (IVT) of mRNAs using the HiScribe™T7 mRNA Kit with CleanCap® Reagent AG (NEB). Cy3-UTP and Cy5-UTP were used replacing 25% of standard UTP in some experiments to fluorescently label the mRNA transcripts. After DNase I treatment, Monarch® RNA Cleanup Kit (NEB) was used to purify RNAs. RNA was analyzed by gel electrophoresis and fragment bioanalyzer (Cornell BRC).

### Mouse experiments

#### Mice

C57BL/6J, BALB/c and Ai14 mice were purchased from Jackson Laboratory. The animals are maintained in a vivarium housing facility by the Center for Animal Resources and Education (CARE) at Cornell University. All animal procedures were approved by the Institutional Animal Care and Use Committee (IACUC) (protocol #2020-0037) and all experiments were performed in accordance with IACUC guidelines for animal care.

#### Systemic administration

Control and Arc EV samples, produced from donor cell culture under sterile conditions, were injected intravenously (retro-orbitally) into mice. EVs were first quantified using epifluorescence intensity reading or NTA, before being diluted in sterile endotoxin-free Dulbecco’s PBS (1X) (w/o Ca++ & Mg++), to ensure the administration of the same amount per kg weight of each mouse. Up to 150 μL of EVs in DPBS was injected into each adult mouse, with 3E+13 total EVs per kg body weight, produced via 1mg cargo mRNA transfected into donor cells. We sometimes store EVs at −80°C (for less than 1 month before injection), adding EDTA as an anticoagulant and sucrose as a cryo-protectant. Mice were carefully monitored daily after injection.

#### Transcardial perfusion

Animals were deeply anaesthetized with Fatal-Plus at a dose of 90 mg/kg and transcardially perfused with 20 ml of ice cold DPBS (1X), followed by 20 ml of 4% paraformaldehyde (PFA) solution. Organs (brain, liver, kidney, spleen, lung, and heart) were quickly extracted and incubated in 4% PFA at 4°C for 24 hrs before being washed and imaged. For cryo-sectioning, brains were transferred to 30% sucrose in PBS solution and allowed to equilibrate for 2 days before being mounted on a freezing microtome using OCT and sectioned coronally into 50 µm thick sections.

### Microscopy

#### Epifluorescence live-cell real-time imaging

Donor and recipient cells were plated in 6-well, 24-well or 96-well plates with phenol-red free medium to be imaged by Cytation 7 cell imager (BioTek), at different time points during transfection, EV production or transduction. Bright field images, together with DAPI, GFP, Cy3 and/or Cy5 channels were often imaged. Total cell (nuclei) numbers were counted based on the DAPI signal using Cytation 7’s own imaging analysis program, with which we can accurately calculate the proportion of total DAPI+ cells showing higher levels of GFP/Cy3/Cy5 fluorescence than the negative controls. In each experiment, we include four replicates (wells) of cells for each sample group, and image the whole well (3 × 3 = 9 tiles for 96-well with a 10X objective for example) that contains ∼20, 000 cells. We also perform serial dilutions of EVs transduced into recipient cells for accuracy of quantification. In addition to the dose of EVs, we also optimize DNA/RNA transfection components in the donor cell culture. More details can be found in supplementary figures.

#### Confocal microscopy

PFA-fixed 50-μm thin cryo-sections or SDS-cleared 200-μm thick tissues were stained via IHC using antibodies, including NeuN-AlexaFluor647 (Abcam, ab190565), GFP-AlexaFluor488 (Invitrogen, A-21311), Iba1 primary antibody rabbit monoclonal (Abcam, ab178846), and Goat anti Rabbit cross absorbed AlexaFluor 546 secondary antibody (Invitrogen, A-11010). Mounting was using ProLong™Gold Antifade Mountant with DAPI (Invitrogen, P36931) for thick sections or PVA DABCO for think sections. CLSM images were acquired using a Zeiss LSM 800 confocal scanning laser microscope with 5X and 20X air objectives.

#### IVIS (in vivo imaging system)

We perform transcardiac perfusion before extracting and fixing organs to be imaged for fluorescence using the IVIS machine. We use the negative controls for background subtraction and normalization, as whole organs display strong autofluorescence. We also applied spectrum unmixing when more than one fluorescence was imaged. We used organs directly injected with fluorophores to optimize spectrum unmixing.

### Cells and cell culture

#### Stable cell line culture

Human embryonic kidney cells (HEK293T and F), human leukemic monocytes (Thp1), murine leukemic monocytes/macrophages (RAW 264.7, ATCC® No. TIB-71™), and immortalized murine dendritic cells (DC2.4) were purchased from ATCC and stored in liquid nitrogen until use. MDA-MB-231 triple negative breast cancer cells and U87 MG glioblastoma cells were kindly gifted by our collaborator, the Cerione lab at Cornell. Cells were maintained following ATCC’s instructions. Cells were transfected with mRNAs or DNAs via liposomes (Invitrogen Lipofectamine or Takara Xfect) or PEI (Polysciences PEImax) following manufacturers’ instructions. No antibiotics were added to any stable cell line culture, especially not during transfection to ensure highest transfection efficiency. Primary donor cells are difficult to obtain with batch-to-batch variations, and so in many optimization/characterization experiments *in vivo*, we used EVs from RAW264.7, a macrophage cell line established from a tumor in a male BALB/c mouse induced with the Abelson murine leukemia virus, and thus also used BALB/c recipient mice in such experiments.

#### Primary BM GM-CSF/IL-4 cultures

Primary bone marrow cells were extracted from wildtype mice (C57BL/6J) after disinfection with 70% ethanol. The femurs and tibiae were dissected and soaked in HBSS (Hank’s Balanced Salt Solution, Gibco) or RPMI-1640 medium (ATCC) supplemented with 1% FBS (Gibco). Both ends of the bone were cut open with surgical scissors, and a 25G needle (of a 20-mL syringe) was inserted into the bone cavity to rinse the BM cells out of the femur, whereas a 27G needle was used for the tibia. A total volume of 20mL HBSS or RPMI-1640 medium was used to slowly (dropwise) flush out BM cells from each femur and a total volume of 10mL was used for each tibia. The cell suspension in the dish was collected and centrifuged at 300X g for 10 min, and the supernatant was discarded. The cell pellet was resuspended with red blood cell (RBC) lysis buffer to lyse the RBCs. Following the second centrifugation, the supernatant was discarded, and the pelleted cells were washed with 1X HBSS and collected. 15 × 10^6^ collected cells were cultured in each tissue-culture-treated T75 flask in 15 ml of complete medium (ATCC RPMI-1640 supplemented with 2-mercaptoethanol (Invitrogen), 10% fetal bovine serum (Gibco), GM-CSF (20 ng/ml, Peprotech) and IL-4 (5 ng/ml, Peprotech).

Half of the medium was removed at day 2 and new warmed medium supplemented with 40 ng/ml GM-CSF and 5 ng/ml IL-4. The culture medium was entirely discarded at day 3 and replaced by fresh warmed medium with 20 ng/ml GM-CSF and 5 ng/ml IL-4. On day 7, all cells in the culture including adherent cells and suspension cells in the culture supernatant, and loosely attached cells harvested by gentle washing with warm PBS were pooled and used as the source for production of leukocyte EVs.

### Donor cell production of EVs

Donor cells were transfected for eraEV production followed by EV isolation and characterization. A typical production time is ∼40 hours after DNA transfection and 16 hours after RNA transfection. Liposome/PEI mediated transfection of DNA vectors (Arc) take place when donor cells reach 60% confluency, followed by the transfection of mRNA transcripts (stabilizer and cargo mRNAs). For DNA transfection, we use Takara’s Xfect system, and for RNA transcription, we use lipofectamine MessengerMAX or PEImax, following manufacturers’ manuals. After transfection, we collect the supernatant from all control and sample groups and purify EVs. Extensive DNase and RNase treatments are essential to ensure low background in the control groups. Based on extensive titration experiments using a wide range of DNA/RNA concentrations, 1250 ng of Arc DNA plasmid or 12 pmol *Arc* mRNA and 8 pmol *cargo* mRNAs are the optimized amount to transfect into each two million of donor cells. For co-transfection of combined cargoes, 4 pmol *cargo* and 4 pmol *stabilizer* were added. We ensure the same amounts of cargo mRNAs are added to each control and experimental group. The lipofectamine MessengerMAX is more efficient transfecting the donor cells but liposomes carrying mRNAs are harder to remove by RNase in the control group leading to a higher background. Transfection always takes place in serum free medium (Expi293 or optiMEM). Not all cargoes are fluorescent for easy visualization, and so we often use the stabilizer A5U with a reporter tail, GFP, to monitor the transfection efficacy in real time. At the end of transfection, we always measure the donor cell number and viability to ensure physiological conditions for EV production.

### Isolation of EVs

Ultrafiltration or density gradient ultracentrifugation was used to isolate EVs. Supernatant cell culture medium was collected from the donor cell cultures and centrifuged for 5 minutes at 300 X g to remove cells and debris. The medium was then passed through 0.45 μm filters to remove larger particles before being concentrated in protein concentrator ultrafiltration tubes with a molecular weight cutoff at 100kDa (MilliporeSigma Amicon Ultra). Endotoxin-free sterile 1X DPBS was used to wash EVs in ultrafiltration tubes. Concentrated EV samples were collected, aliquoted, with some stained by a plasma membrane dye and/or an anti-Arc antibody. After the staining, excessive washes using 1X DPBS remove unbound free dyes and antibodies. 1X DPBS without EV was used as a negative control, stained, and washed to the same degree as the experimental groups. With one more extra wash after the no-EV control turned colorless, we would finish washing all samples. For density gradient centrifugation, we add a layer of 60% sucrose cushion on the bottom, a layer of 30% iodixanol in the middle and a layer of 15% sucrose and then supernatant medium on top. Alternating two materials is for better separation of layers and Arc EVs settle in the 30% iodixanol layer, which is non-toxic to cells. The supernatant medium is generally concentrated for 100X for each adult recipient mouse, and for 20X for *in vitro* recipient cell culture. This is necessary due to a big loss during the EV production (high-level uptake by donor cells), collection and storage (EV aggregation despite the usage of anticoagulant and cryoprotectant).

### Fluorescent NTA

The NanoSight NS300 instrument (NanoSight Ltd.) allows the detection of nanovesicles labeled with stable fluorophores. It uses a 405 nm (violet) laser diode to excite suitable fluorophores whose fluorescence can then be determined using a matched 430 nm long-pass filter. Measurements are made in fluorescence mode with the long-pass filter in place so that only fluorescently emitted light is measured. The fluorescent particles are individually tracked in real time, from which labeled particle size and concentration can be determined. Under light scatter mode, the total number of particles can be measured and subsequently compared to the concentration of labeled particles. EVs were labeled, by fluorescent antibodies against general EV marker (CD63), Arc, Syncytin A or a green plasma membrane dye for NTA measurements of CD63+ EV number, Arc EV proportion, and total EV number, respectively.

### Negative stain electron microscopy (NSEM)

For all negative stain specimens, copper 200-mesh grids coated with Formvar and carbon (Electron Microscopy Sciences or Ted Pella, Redding, CA) were glow discharged for 20–45 s in a vacuum chamber at 30mA. 3.5 μL sample was then applied to the grid for 35–45 s and excess sample was wicked away using filter paper. Grids were then immediately washed 2–4 × for 5 s with 30 μL water droplets, then once with 1% uranyl acetate (UA) on parafilm. Excess water/UA was wicked away and then a final droplet of UA was applied for 30 s. Excess UA was wicked away and grids were air dried for 30–60 s. Imaging was performed using either an FEI T12, FEI Tecnai Spirit microscope operated at 120 kV equipped with a Gatan Orius SC200B CCD camera or JEOL 1400 electron microscope.

### Cryogenic electron microscopy (Cryo-EM)

Degassed 2/2-3C C-flat grids (Electron Microscopy Sciences, Hatfield, PA) were glow discharged for 45 s at 30 mA. Sample was applied to the grid 2 times for 30 s, and the grid was plunge frozen in liquid ethane using a FEI Vitrobot Mark IV. Micrographs were acquired using a FEI Tecnai G^2^ F20 microscope operated at 200 kV, equipped with a FEI Falcon II direct detector. The nominal defocus was 1.3 μm.

### Arc protein capsid assembly

*GFP* and *A5U-GFP* mRNAs were added to purified rat Arc (RNA−) (2 mg/mL in low salt buffer: 50 mM NaCl, 50 mM Tris, pH 8 at RT) at a nucleic acid:protein ratio of 21.6% (w/w) (corresponding to 1 molecule of Arc to 30 nucleotides). Reactions were then dialyzed into the assembly buffer (1 M NaCl, 50 mM Tris, pH 7.4 with 0.5 mM EDTA) 24-48 hrs in the cold room before being analyzed by DLS, NTA and AFM (atomic force microscopy).

### Flow Cytometry Analysis

Cells were stained in ice-cold DPBS containing FCS (2%) and EDTA (2 mM) using appropriate antibody-fluorophore conjugates. Murine FcR blocking reagent (Miltenyi Biotec) was used to block unwanted binding of antibodies to mouse cells expressing Fc receptors. Multiparameter analysis was performed and analyzed on a Attune analyzer (Invitrogen). The following antibodies were purchased from BD Bioscience: anti-Ly6c (clone AL-21), anti-CD45.1 (clone A20), anti-B220 (clone RA3-6B2); from Biolegend: anti-F4/80 (clone C1:A3-1), CD11c (clone N418); from eBioscience: anti-MHCII I-A/I-E (clone M5/114.15.2), anti-CD11b (cloneM1/70), anti-CD14 (clone Sa2-8), and anti-Syncytin A (thermofisher **#** BS-2962R). Corresponding isotype-matched irrelevant specificity controls were purchased from BD, Biolegend, and eBioscience.

### RNA immunoprecipitation followed by real time quantitative PCR

5 × 10^6^ donor cells and EVs from the supernatant medium were harvested by centrifugation and ultrafiltration respectively. After cell and EV lysis, an Arc antibody (Synaptic systems 156 003) was used to immunoprecipitate the Arc capsids carrying the mRNA cargo inside. Homogenates were pelleted at 200×***g*** for 5 min at 4°C to remove tissue debris. Supernatants were removed, diluted from 2 mL to 4 mL, and rocked at 4°C for 10 min before being pelleted at 17,000×***g*** for 10 min at 4°C to remove insoluble material. Cleared supernatants were removed, a small aliquot was taken as the input, and the remainder used for immunoprecipitation. Supernatants were immunoprecipitated with either Arc antibody (rabbit polyclonal, custom-made; Protein Tech) or normal rabbit IgG (Santa Cruz Biotechnology, Santa Cruz, CA) at 1 μg/500 μL lysate for 2 h at 4°C with gentle rocking. Following antibody incubation, a 10% volume of washed 50/50 Protein A bead slurry (Thermo Fisher Scientific) was added to the antibody/lysate mixture and incubated for an additional hour at 4°C with rocking. Bead-antibody complexes were then pelleted briefly at low speed, supernatants were removed, and beads were washed three times with IP buffer. Washed beads were then resuspended in 200 μL IP buffer. With half of the bead slurry, protein was eluted from the beads with 17 μL 4X Laemlli buffer for 5 min at RT, then 50 μL IP buffer was added and the solution was removed from the beads into a new tube and heated at 70°C for 5 min. The input (10% lysate volume) and 30 μL each of the IgG and antibody elutions were separated by SDS-PAGE on a 10% acrylamide gel and immunoblotted as described above. The bands for the input and IgG and Arc elutions were analyzed using the Gel Analysis plugin in ImageJ, and the data were represented graphically as a ratio of the signal from each elution over the input signal from each individual mouse. With the other half of the bead slurry, the IP buffer was adjusted to 1% SDS and 0.8 mg Proteinase K (New England Biolabs, Ipswich, MA) was added. Samples were then incubated at RT for 30 min with rocking and total RNA was extracted as described. For all samples, total RNA was extracted using TRIzol (Thermo Fisher Scientific). TRIzol-extracted samples were mixed 5:1 with chloroform, incubated at RT for 3 min, and pelleted at 12,000×***g*** at 4°C for 10 min. The resulting aqueous phase was taken and mixed 1:1 with isopropanol, incubated at RT, and pelleted at 12,000×***g*** at 4°C for 10 min. The resulting supernatant was removed, and pellet washed with cold 75% ethanol. Washed pellets were then repelleted at 7500×***g*** for 5 min at 4°C. The supernatant was removed and dried pellets were resuspended in ddH_2_O. RNA was then reverse transcribed into cDNA and analyzed by RT-qPCR. Total RNA concentrations were measured by A_260/280_ on a Nanodrop (Thermo Scientific). Reverse transcription reactions were carried out using a High-Capacity cDNA Reverse Transcription Kit (Applied Biosystems, Foster City, CA) with 100–200 ng of RNA as template. Resulting cDNA was prepared for qPCR using PowerUp SYBRgreen Master Mix (Thermo Fisher Scientific) in a 96-well plate with primers against:

**Table.**
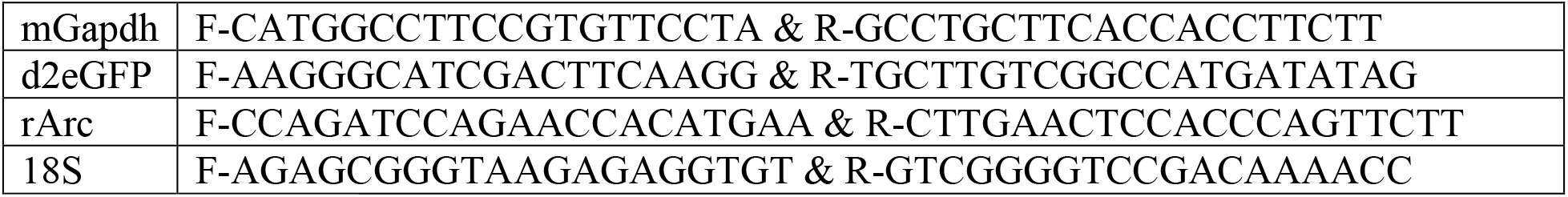
qPCR primers.

qPCR was performed on a BioRad CFX Real Time PCR System using the following protocol: Pre-incubation: 50°C for 2 min, 95°C for 2 min. Amplification: 40 cycles of 95°C for 15 s, 60°C for 15 s, and 72°C for 1 min. Melt curve: 95°C for 1 s, 60°C for 20 s, continuous ramp at 0.1 5°C/s up to 95°C. Ct values of greater than 30 were considered undetectable. Differences in expression were determined using the standard curve method, where a standard DNA sample was serially diluted (10-fold), analyzed for the gene of interest, and the linear equation calculated. The resulting linear equation was used to determine where the Ct values of test samples fell within the standard curve and the result was transformed (log_10_) to reflect the dilution of the standard sample. Differences were calculated measuring the fold-change from the average of the control values for any given group (test/average control).

### Statistical analysis

All data are reported as mean ± SD, dot plots, or bar graphs. Student T test, one-way or two-way ANOVA were used to compare differences among multiple groups. All analyses were carried out with GraphPad Prism 9 Software. The criterion for statistical significance was *P* < 0.05. For *in vitro* experiments, millions of cells were imaged with at least five regions selected and further analyzed. Tens of thousands of cells in each field of view were quantified for mean fluorescence intensity, integrated density and/or percentage of cells showing signals above defined thresholds subtracting the background levels based on the negative control and sometimes FMO (fluorescence minus one) controls. Error bars were calculated from such independent fields of view. All experiments were repeated for at least three times with results presented when showing the same trend. For *in vivo* experiments, we conducted a power analysis^39^ examining predicted statistical power using various cohort sizes and difference in mRNA accumulation in the brain, assuming approximately normally distributed residuals, a correlation of 0.5, and an alpha of 0.05. Based on this analysis, we intend to use 5 mice for each control and experimental group, which should give a statistical power of at least 0.9 if we observe at least 40% difference in the accumulation of fluorescent mRNA cargoes in the brain between sCargo+/Arc‒ control and eraEV groups. This condition is very stringent, as there is endogenous Arc expressing in the donor cells or animals, this group already show a smaller but significant increase in the level of neuronal cargo mRNAs compared to other negative control groups. For either variety of experiment, data will be considered statistically meaningful at a p value < 0.05.

### ImageJ macro-code for batch image quantification

<u>Cell count overlapping bare outlines</u> (Parameters in this code may be changed accordingly depending on the raw images):

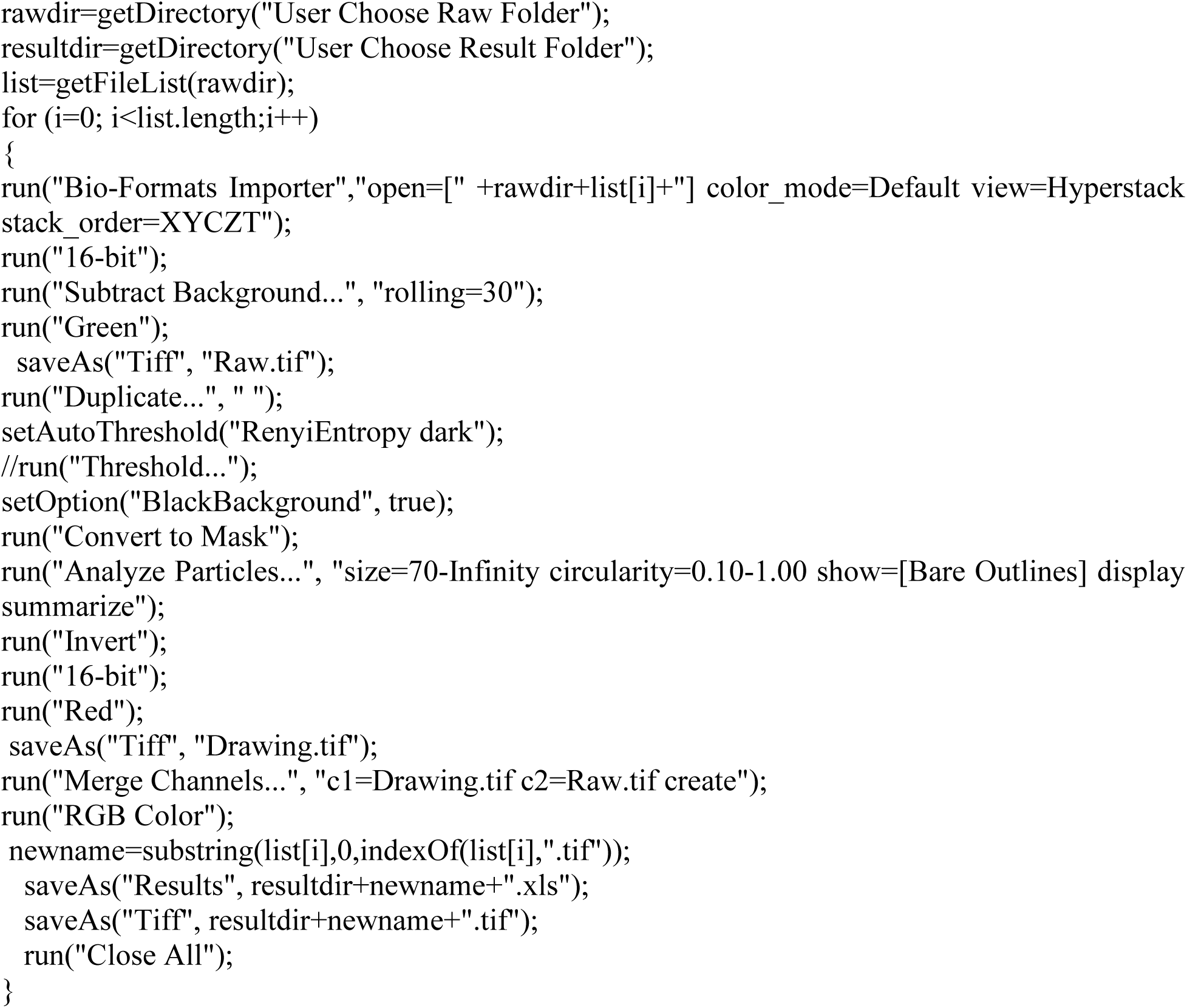

## Acknowledgments

We thank Rebecca Williams, Tina Abratte and the BRC Imaging Facility at the Cornell Institute of Biotechnology (RRID:SCR_021741) for imaging experiments, with NIH S10OD025049 for the IVIS-Spectrum optical imager in Cornell’s BRC Imaging Facility. We thank P. Schweitzer and the BRC Genomics Facility (RRID:SCR_021727) at the Cornell Institute of Biotechnology for sequencing experiments. We thank Lydia Tesfa, Jaclyn Mahoney and the BRC Flow Cytometry Facility (RRID:SCR_021740) for flow cytometry data. We thank Trevor Totman, Erica Feldman, Faith Burgus, and the CARE at Cornell for their service and advice caring for animals. We thank Robert Felt and the IACUC, Cornell for their help composing and managing animal protocols. We thank Andrew Recknagel for his valuable advice on imaging. S.J. acknowledges start-up support from Cornell University, including Robert Langer ‘70 Family and Friends Professorship and Cornell NEXT Nano Initiative.

## Author contributions

Conceptualization: Wenchao Gu, Robert Langer, Shaoyi Jiang.

Data acquisition: Wenchao Gu, Sijin Luozhong, Simian Cai, Ketaki Londhe, Nadine Elkasri, Robert Hawkins, Margaret Cruz, Andy Chang, Wenting Gao, Tara Sarmiento.

Analysis and interpretation of results: Wenchao Gu.

Advice, support, and supervision: Wenchao Gu, Zhefan Yuan, Kai Su-Greene, Patrick McMullen, Chunyan Wu, Changwoo Seo, Akash Guru, Chris Schaffer, Nozomi Nishimura, Richard Cerione, Melissa Warden, Robert Langer, Shaoyi Jiang.

Writing (original draft): Wenchao Gu.

Writing (review and editing): Chris Schaffer, Nozomi Nishimura, Richard Cerione, Melissa Warden, Robert Langer, Shaoyi Jiang.

## Data and materials availability

All data are available in the main text or the supplementary materials.

## References

1 Dong, X. Current Strategies for Brain Drug Delivery. Theranostics 8, 1481–1493 (2018). https://doi.org:10.7150/thno.21254

2 Pardridge, W. M. Blood-Brain Barrier and Delivery of Protein and Gene Therapeutics to Brain. Front Aging Neurosci 11, 373 (2019). https://doi.org:10.3389/fnagi.2019.00373

3 Sweeney, M. D., Zhao, Z., Montagne, A., Nelson, A. R. & Zlokovic, B. V. Blood-Brain Barrier: From Physiology to Disease and Back. Physiol Rev 99, 21–78 (2019). https://doi.org:10.1152/physrev.00050.2017

4 Varatharaj, A. & Galea, I. The blood-brain barrier in systemic inflammation. Brain Behav Immun 60, 1–12 (2017). https://doi.org:10.1016/j.bbi.2016.03.010

5 Nian, K., Harding, I. C., Herman, I. M. & Ebong, E. E. Blood-Brain Barrier Damage in Ischemic Stroke and Its Regulation by Endothelial Mechanotransduction. Front Physiol 11, 605398 (2020). https://doi.org:10.3389/fphys.2020.605398

6 Banks, W. A. et al. Transport of Extracellular Vesicles across the Blood-Brain Barrier: Brain Pharmacokinetics and Effects of Inflammation. Int J Mol Sci 21 (2020). https://doi.org:10.3390/ijms21124407

7 Yuan, D. F. et al. Macrophage exosomes as natural nanocarriers for protein delivery to inflamed brain. Biomaterials 142, 1–12 (2017). https://doi.org:10.1016/j.biomaterials.2017.07.011

8 Haney, M. J. et al. Exosomes as drug delivery vehicles for Parkinson’s disease therapy. J Control Release 207, 18–30 (2015). https://doi.org:10.1016/j.jconrel.2015.03.033

9 Man, S., Ubogu, E. E. & Ransohoff, R. M. Inflammatory cell migration into the central nervous system: a few new twists on an old tale. Brain Pathol 17, 243–250 (2007). https://doi.org:10.1111/j.1750-3639.2007.00067.x

10 Seguin, R., Biernacki, K., Rotondo, R. L., Prat, A. & Antel, J. P. Regulation and functional effects of monocyte migration across human brain-derived endothelial cells. J Neuropathol Exp Neurol 62, 412–419 (2003). https://doi.org:10.1093/jnen/62.4.412

11 Erickson, M. A. & Banks, W. A. Age-Associated Changes in the Immune System and Blood-Brain Barrier Functions. International Journal of Molecular Sciences 20 (2019). https://doi.org:ARTN163210.3390/ijms20071632

12 Aslan, C. et al. Exosomes for mRNA delivery: a novel biotherapeutic strategy with hurdles and hope. BMC Biotechnol 21, 20 (2021). https://doi.org:10.1186/s12896-021-00683-w

13 Li, M. et al. Analysis of the RNA content of the exosomes derived from blood serum and urine and its potential as biomarkers. Philos Trans R Soc Lond B Biol Sci 369 (2014). https://doi.org:10.1098/rstb.2013.0502

14 Chevillet, J. R. et al. Quantitative and stoichiometric analysis of the microRNA content of exosomes. Proc Natl Acad Sci U S A 111, 14888–14893 (2014). https://doi.org:10.1073/pnas.1408301111

15 Valadi, H. et al. Exosome-mediated transfer of mRNAs and microRNAs is a novel mechanism of genetic exchange between cells. Nat Cell Biol 9, 654–659 (2007). https://doi.org:10.1038/ncb1596

16 Luan, X. et al. Engineering exosomes as refined biological nanoplatforms for drug delivery. Acta Pharmacol Sin 38, 754–763 (2017). https://doi.org:10.1038/aps.2017.12

17 Usman, W. M. et al. Efficient RNA drug delivery using red blood cell extracellular vesicles. Nat Commun 9, 2359 (2018). https://doi.org:10.1038/s41467-018-04791-8

18 Momen-Heravi, F., Bala, S., Bukong, T. & Szabo, G. Exosome-mediated delivery of functionally active miRNA-155 inhibitor to macrophages. Nanomedicine 10, 1517–1527 (2014). https://doi.org:10.1016/j.nano.2014.03.014

19 Wang, J. H. et al. Anti-HER2 scFv-Directed Extracellular Vesicle-Mediated mRNA-Based Gene Delivery Inhibits Growth of HER2-Positive Human Breast Tumor Xenografts by Prodrug Activation. Mol Cancer Ther 17, 1133–1142 (2018). https://doi.org:10.1158/1535-7163.MCT-17-0827

20 Pastuzyn, E. D. et al. The Neuronal Gene Arc Encodes a Repurposed Retrotransposon Gag Protein that Mediates Intercellular RNA Transfer. Cell 172, 275–288 e218 (2018). https://doi.org:10.1016/j.cell.2017.12.024

21 Segel, M. et al. Mammalian retrovirus-like protein PEG10 packages its own mRNA and can be pseudotyped for mRNA delivery. Science 373, 882–889 (2021). https://doi.org:10.1126/science.abg6155

22 Ashley, J. et al. Retrovirus-like Gag Protein Arc1 Binds RNA and Traffics across Synaptic Boutons. Cell 172, 262–274 e211 (2018). https://doi.org:10.1016/j.cell.2017.12.022

23 Comas-Garcia, M., Davis, S. R. & Rein, A. On the Selective Packaging of Genomic RNA by HIV-1. Viruses 8 (2016). https://doi.org:10.3390/v8090246

24 Dynes, J. L. & Steward, O. Arc mRNA docks precisely at the base of individual dendritic spines indicating the existence of a specialized microdomain for synapse-specific mRNA translation. J Comp Neurol 520, 3105–3119 (2012). https://doi.org:10.1002/cne.23073

25 Fila, M., Diaz, L., Szczepanska, J., Pawlowska, E. & Blasiak, J. mRNA Trafficking in the Nervous System: A Key Mechanism of the Involvement of Activity-Regulated Cytoskeleton-Associated Protein (Arc) in Synaptic Plasticity. Neural Plast 2021, 3468795 (2021). https://doi.org:10.1155/2021/3468795

26 Paolantoni, C. et al. Arc 3’ UTR Splicing Leads to Dual and Antagonistic Effects in Fine-Tuning Arc Expression Upon BDNF Signaling. Front Mol Neurosci 11, 145 (2018). https://doi.org:10.3389/fnmol.2018.00145

27 Giorgi, C. et al. The EJC factor eIF4AIII modulates synaptic strength and neuronal protein expression. Cell 130, 179–191 (2007). https://doi.org:10.1016/j.cell.2007.05.028

28 Booth, A. M. et al. Exosomes and HIV Gag bud from endosome-like domains of the T cell plasma membrane. J Cell Biol 172, 923–935 (2006). https://doi.org:10.1083/jcb.200508014

29 Comas-Garcia, M. et al. Dissection of specific binding of HIV-1 Gag to the ‘packaging signal’ in viral RNA. Elife 6 (2017). https://doi.org:10.7554/eLife.27055

30 Brigham, B. S., Kitzrow, J. P., Reyes, J. C., Musier-Forsyth, K. & Munro, J. B. Intrinsic conformational dynamics of the HIV-1 genomic RNA 5’UTR. Proc Natl Acad Sci U S A 116, 10372–10381 (2019). https://doi.org:10.1073/pnas.1902271116

31 Blakemore, R. J. et al. Stability and conformation of the dimeric HIV-1 genomic RNA 5’UTR. Biophys J 120, 4874–4890 (2021). https://doi.org:10.1016/j.bpj.2021.09.017

32 Carlson, L. A., Bai, Y., Keane, S. C., Doudna, J. A. & Hurley, J. H. Reconstitution of selective HIV-1 RNA packaging in vitro by membrane-bound Gag assemblies. Elife 5 (2016). https://doi.org:10.7554/eLife.14663

33 Madisen, L. et al. A robust and high-throughput Cre reporting and characterization system for the whole mouse brain. Nat Neurosci 13, 133–140 (2010). https://doi.org:10.1038/nn.2467

34 Rossler, K. et al. Expression of leucocyte adhesion molecules at the human blood-brain barrier (BBB). J Neurosci Res 31, 365–374 (1992). https://doi.org:10.1002/jnr.490310219

35 Owens, T., Bechmann, I. & Engelhardt, B. Perivascular spaces and the two steps to neuroinflammation. J Neuropathol Exp Neurol 67, 1113–1121 (2008). https://doi.org:10.1097/NEN.0b013e31818f9ca8

36 Zozulya, A. L. et al. Dendritic cell transmigration through brain microvessel endothelium is regulated by MIP-1alpha chemokine and matrix metalloproteinases. J Immunol 178, 520–529 (2007). https://doi.org:10.4049/jimmunol.178.1.520

37 Larochelle, C., Alvarez, J. I. & Prat, A. How do immune cells overcome the blood-brain barrier in multiple sclerosis? FEBS Lett 585, 3770–3780 (2011). https://doi.org:10.1016/j.febslet.2011.04.066

38 Helft, J. et al. GM-CSF Mouse Bone Marrow Cultures Comprise a Heterogeneous Population of CD11c(+)MHCII(+) Macrophages and Dendritic Cells. Immunity 42, 1197–1211 (2015). https://doi.org:10.1016/j.immuni.2015.05.018

39 Cohen, J. The statistical power of abnormal-social psychological research: a review. J Abnorm Soc Psychol 65, 145–153 (1962). https://doi.org:10.1037/h0045186

