## Supplemental materials for "Engineered retrovirus-like nanocarriers for messenger RNA delivery into neurons"

**This session includes:**

Supplementary Discussion

Figs. S1 to S10

Tables S1

References (1-2)

**Supplementary Discussion**

We optimized EV production conditions and characterization methods. To produce Arc EVs, we used the rat Arc capsid for easier characterization, as it differs from endogenous Arc in human or mouse cell lines and *in vivo* mouse models. Extensive and comprehensive titration and time lapse experiments were carried out to optimize EV production (Fig. S1-2 and data not shown). A variety of nucleic acid transfection methods were tested in each donor cell type, including PEI-mediated methods and a variety of liposome-mediated methods (Fig. S1-2). Each method possesses different transfection efficacy in different donor cells. For example, PEI is effective in HEK293 but is not efficient transfecting RAW 264.7 macrophages. We often include a reporter in our RNA stabilizer to monitor the transfection efficacy in real time. We aim to tune the fine balance between donor cell viability (ensure high EV viability, Fig.S1) and effective transfection (ensure high quantity of engineered EVs, Fig. S2).

Our first key result is that A5U is required for either the production or preservation of Arc EVs. To characterize Arc+ EV subpopulation among total EVs, we first performed fluorescent NTA with a green laser, to capture Arc+ EVs labeled by antiArc-Alexa532 (EV membrane permeabilized with Triton X-100). When applying a reduced camera level, with which the non-stained negative control EVs show no fluorescence, only Arc+ EVs were analyzed and quantified. When the camera level was adjusted to the maximum intensity, all EVs became visible due to light scattering by particles, including the unlabeled non-fluorescent populations. When fluorescent antibodies against general EV markers, CD63, CD9 and CD81, were used to label and identify total EVs, we noticed an increase in the size measured via NTA. This also applies to other fluorescent plasma membrane dyes (e.g., CellMask), bound the outer membrane of EVs. Arc, on the other hand, is intra-vesicular and we permeabilized the EV membrane with a detergent, Triton X-100, allowing the antibody to penetrate and label the capsid. Indeed, the mean size of non-labeled EVs from the Arc+ sample group measures almost the same as the fluorescently labeled vesicles (slightly larger likely due to a density change), indicating that intra-vesicular anti-Arc fluorescent labelling of Arc+ EVs does not increase their size measured by NTA. Moreover, increasing recent evidence suggest that the classical general EV markers are biased towards different EV subpopulations. For example, CD63 has been found at higher levels in exosomes, whereas CD9/81 may mostly mark ectosomes[1].

Our next key finding is that Arc promotes mRNA loading into EVs and A5U further improves mRNA package into the Arc EV. With fluorescent mRNA *in vitro* transcribed by incorporating Cy3-UTPs, we could visualize cargo loading from donor cells and cargo uptake by recipient cells. We did liposome mediated RNA transfection to deliver the cargo mRNAs. There could be potential fusion of liposome-mRNAs with Arc EVs to provide false positive signals, but this shall be a rare event, due to the following reasons: (a) At 4-6 hours after the transfection, excessive liposome/mRNAs were removed, by rinsing the donor cells twice with fresh warm medium before a medium change. The donor cell culture was imaged 40 hours later, when no fluorescent liposomes shall remain. This was proven by the lack of fluorescent extracellular particles in Arc– control groups, which had the same amount of liposome-$_{Cy3}$RNA transfected as in the Arc+ groups. In fact, the removal of liposome-RNA is a critical step in the Xfect (RNA) protocol from Takara. (b) Moreover, Arc capsids assemble at the plasma membrane and bud out of donor cells via direct outward budding, which is a tightly regulated and coordinated process taking place in a distinct subcellular location from where transfected liposomes would accumulate (endosome, MVB and lysosome). Furthermore, we have cargo only control group with the same amount of liposome-cargo loaded, in addition to the mock transfection control, the capsid only control, the cargo w/o stabilizer control, and the capsid w/ cargo w/o stabilizer control. None of the control groups showed cargo encapsulation as efficient and stable as the experimental group (capsid+/stabilizer+/cargo+).

The efficiency and dynamics of cargo loading depend on the fine balance between the amount of capsid and cargo components transfected (Fig. S5). Excessive *Arc* increases cargo loading capacity while reducing specificity (Fig.S5A-B). A low capsid:cargo ratio enabled preferential loading of *A5U+ mRNAs,* among other abundant RNAs, including overexpressed *rArc*, a cytosolic housekeeping gene (*GAPDH*), and *18S*, one of the most prominent RNA species found in ectosomes[2] (Fig. S5B). However, this ratio does not give rise to a sufficient proportion of engineered EVs. We eventually determined the most effective method for EV production from the donor cell culture being a DNA transfection of the pCMV-Arc vector first to accumulate Arc proteins for 24 hours, followed by an RNA transfection of the stabilizer and cargoes.

Our next key result is that Leukocyte derived eraEVs efficiently deliver mRNA across the BBB specifically targeting neuro-inflammation. Here we provide more information about the assessment of EV production from primary donor cells in real time: No matter which cargo to load and deliver, we always use A5U to stabilize the Arc capsid, adding a reporter tail to monitor EV production rate in real time. Depending on the confluency of donor cells, the time it takes to reach the maximum transfection efficacy can vary (Fig. S7C1-C2). Despite our best effort to keep donor cell culture consistent, these primary cells do vary from batch to batch. We thus monitored GFP expression in donor cells via live cell imaging, aiming to collect EVs at their peak of yield, typically ~40 hours post transfection. At 40 hours, the quantification of total EVs by CMDR epifluorescence also confirmed the comparable level of EV secretion among sample groups (Fig. S7D, red). Similar to when EVs are generated in human cell lines, Arc+/A5U+ preserved the highest proportion of Arc EVs among total EVs from primary donor BM-DC/M (Fig. S7D, green).

### Supplementary Figures

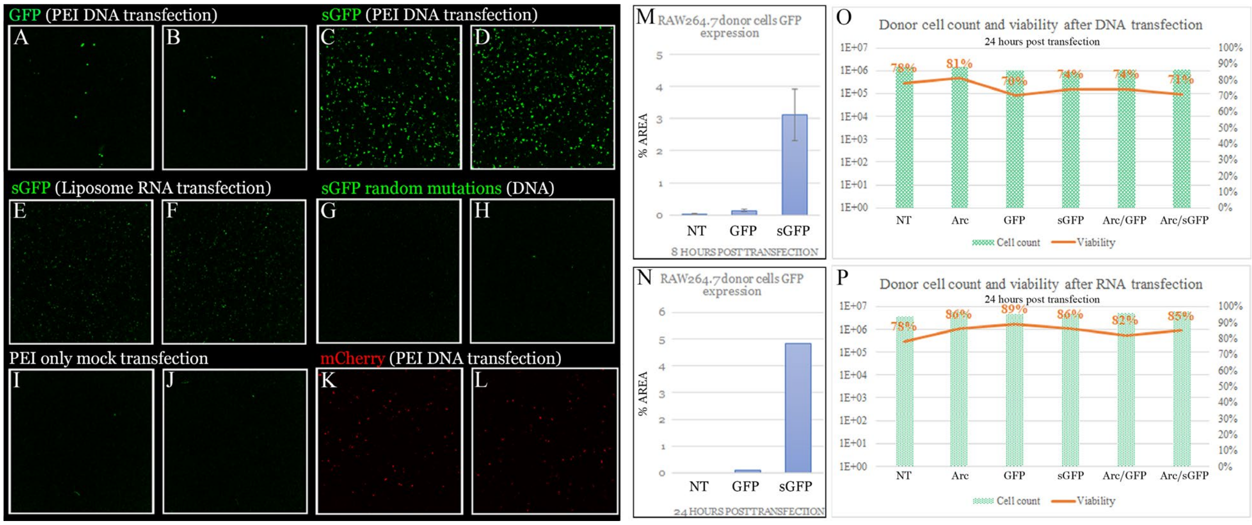


**Fig. S1. Transfection of engineered DNA constructs and RNA transcripts into donor cells to test their functionality.** DNA constructs (A-D) and RNA transcripts (E-F) to encode the cargo have been verified in various donor cells with a radon mutation negative control (G-H), a mock transfection control (I-J), and a distinct fluorophore mCherry control (K-L), with live-cell epifluorescence imaging applied to monitor expression in real time lapse (M-N). **(O-P)** The number and viability of donor cells at the end of EV production was always recorded and compared to ensure a good and comparable quality of EVs produced among control and experimental groups. (O) is from an DNA transfection experiment, whereas (P) is from an RNA transfection experiment.

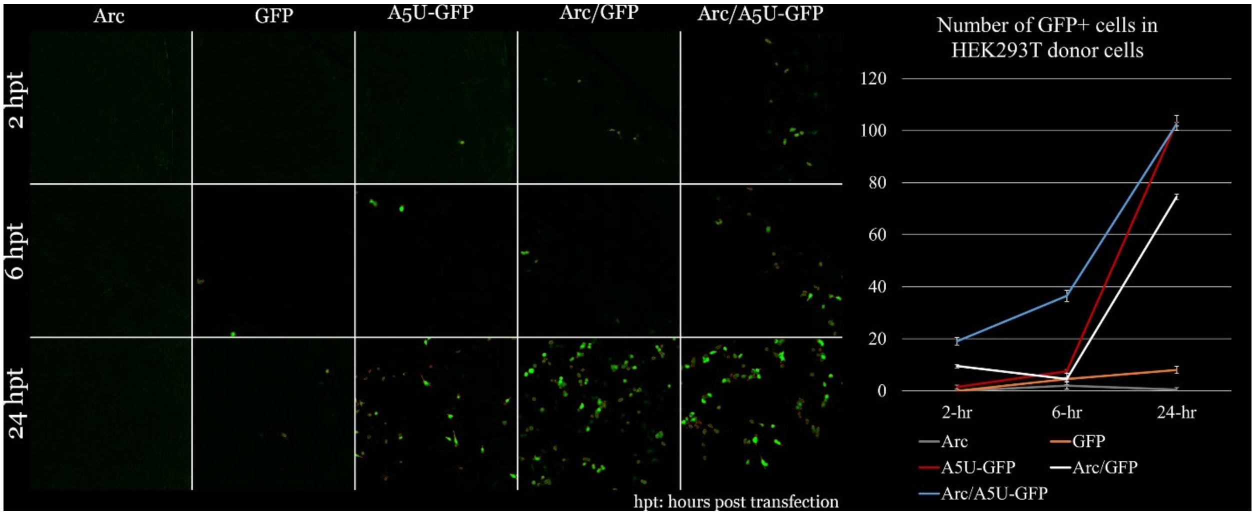


**Fig. S2. DNA transfection time lapse with quantification of GFP expression.** PEI-DNA transfection into HEK293 cells provides DNA plasmid templates, which have longer half-lives than RNA transcripts from RNA transfection, maintaining long-lasting production of both capsid and cargo mRNAs and proteins, we observed a significant and stable increase in GFP expression by transfecting the capsid construct.

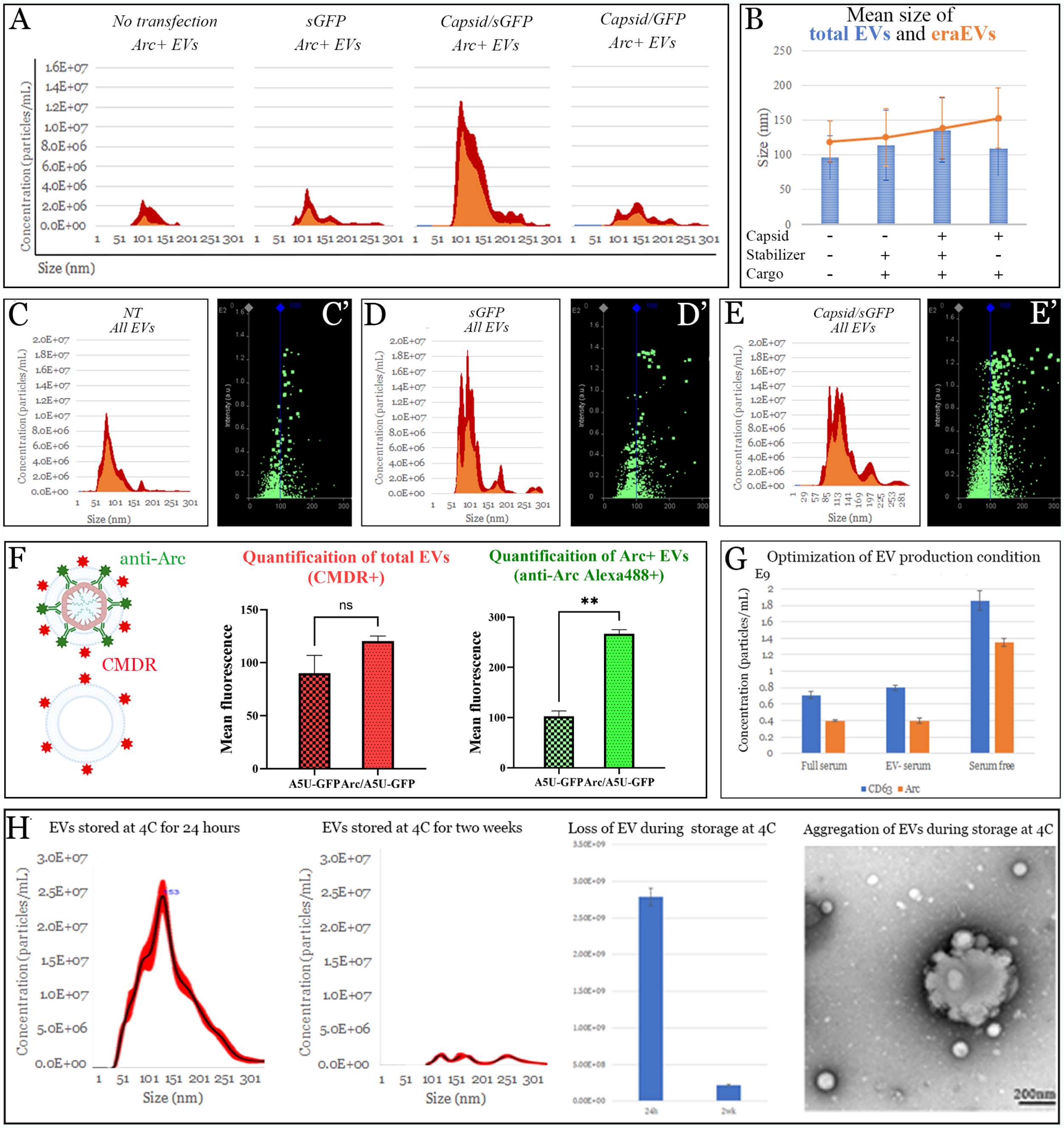


**Fig. S3. EV characterization and EV production optimization. (A)** Concentrations of particles are plotted as a function of their sizes. Arc-EVs appear larger than non-Arc-EVs. Only with the stabilizer A5U motif, a significant amount of capsid+ EVs were counted. **(B)** The mean size of total EVs matches that of capsid+ eraEVs in the sample group with the stabilizer, whereas the other groups have smaller mean sizes due to the presence of a larger proportion of capsid- EVs. **(C-E)** NTA results suggested that the production of total EVs was comparable among all transfected control and experimental groups. EVs were produced for 48 hours before collection. **(C'-E')** An increased number of larger (or denser) vesicles were produced by *Arc* transfection. **(F)** Epi-fluorescence of total EVs (CMDR+) and Arc+ EVs was measured to quantify EV subpopulations, showing the same trend as fluorescent NTA as in Fig.1F. **(G)** Serum free production of EVs led to a higher-level secretion of mouse CD63+ EVs and a larger proportion of eraEVs among all EVs secreted. **(H)** EVs can be stored at 4°C for a short period of time (e.g., 24 hours) but aggregate rapidly.

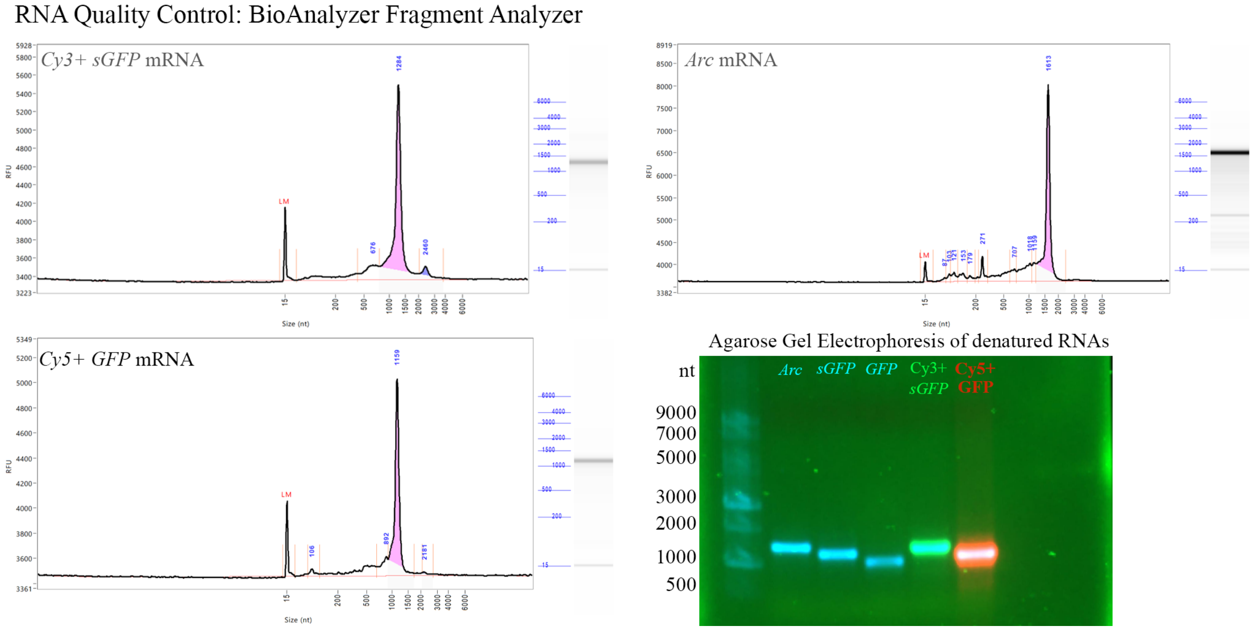


**Fig. S4. The RNA quality control (QC).** BioAnalyzer fragment analysis of *in vitro* synthesized mRNA transcripts showed a good purity and quality as well as the correct sizes of both fluorescent and non-fluorescent RNAs. These RNA aliquots were stored at -80°C for six-month and then 24 hours at 4°C after defrosting before this analysis. The gel electrophoresis was done immediately after RNA synthesis and purification.

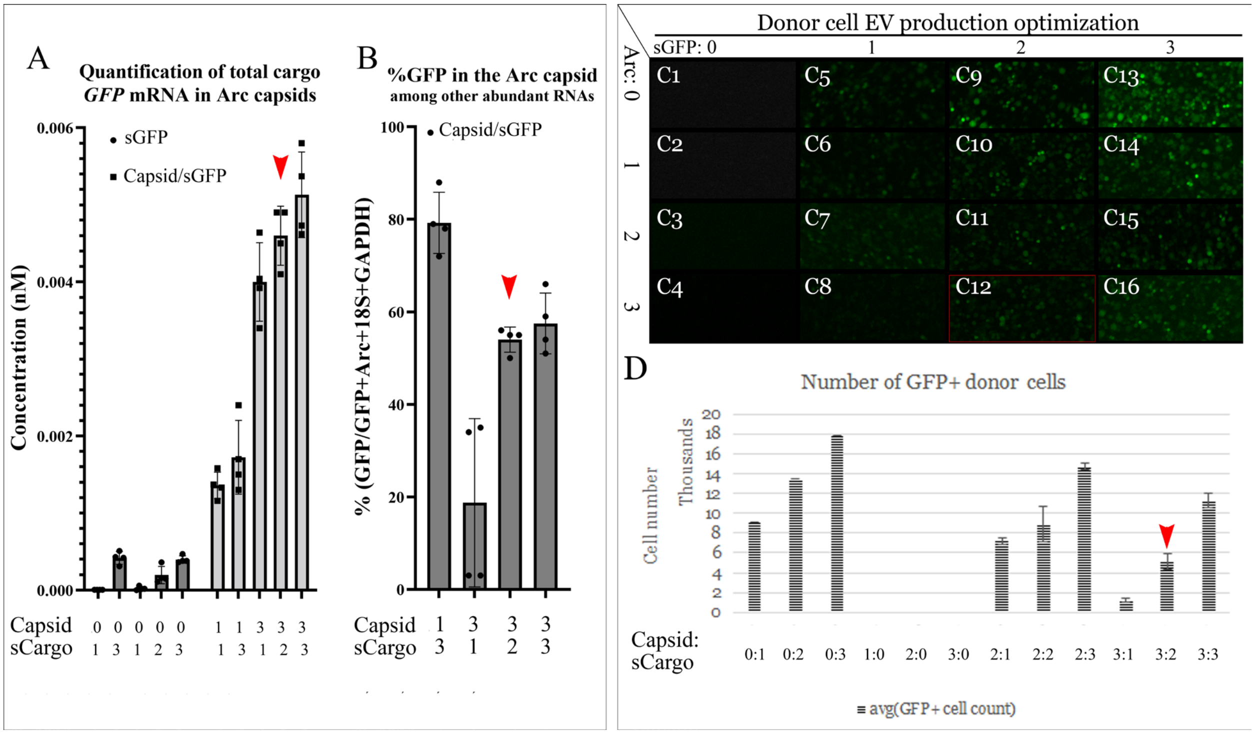


**Fig. S5. Optimization of the ratio between capsid and cargo mRNA transfection components.** Different molar ratios between capsid and cargo mRNAs were transfected into RAW264.7 cells. *Capsid:Cargo*=1:1 is when 1.54 µg *Arc* and 0.93 µg *GFP or* 1.13 µg *A5U-GFP* were transfected into each 2,000,000 cells in a T25 flask. **(A)** Cargo mRNA was loaded more efficiently with an increase in capsid *Arc* mRNA transfected. **(B)** However, an increased amount of other abundant cellular RNAs were also loaded, reducing the purity of cargo. A fine balance between efficient cargo loading and cargo selectivity is critical for the system's functionality. For example, *Capsid:Cargo*=1:3 resulted in inefficient encapsulation of cargo and therefore did not transfer cargo well into recipient cells, whereas *Capsid:Cargo*=3:1 loaded a large amount of excessive $_{capsid}$*Arc* mRNA. **(C)** An increase in the amount of $_{capsid}$*Arc* mRNA led to decreased $_{cargo}$GFP protein expression initially but eventually became comparable to the level when the cargo was transfected alone. The quantification of GFP+ donor cells is shown in **(D)**.

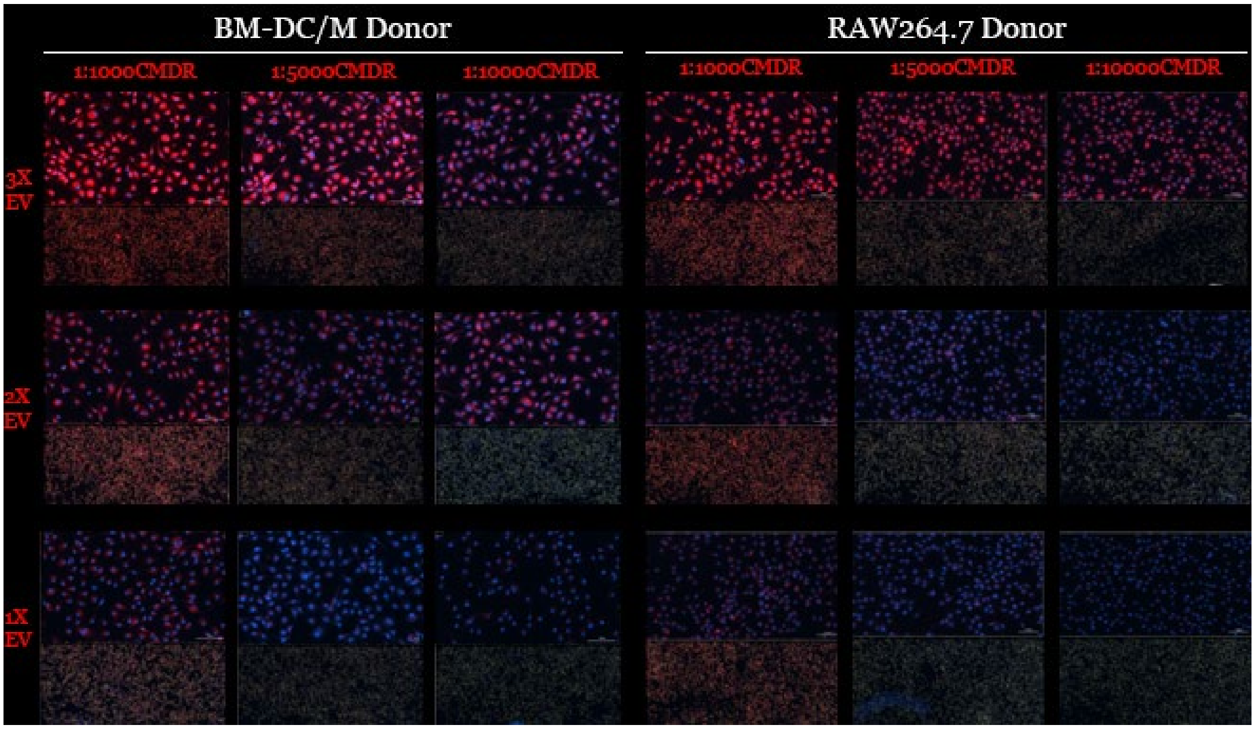


**Fig. S6. CMDR dye optimization for the quantification of total EV uptake.** Staining with a plasma membrane stain, CMDR, was carried out to facilitate the quantification of EVs in solution and in recipient cells. The number of EVs to be transferred to recipient cells was also optimized. We eventually decided on the 1:5000 CMDR dye concentration and the 2X EV concentration, meaning that EVs collected from 2 donor cells producing for 24-40 hours to be used for 1 recipient cell.

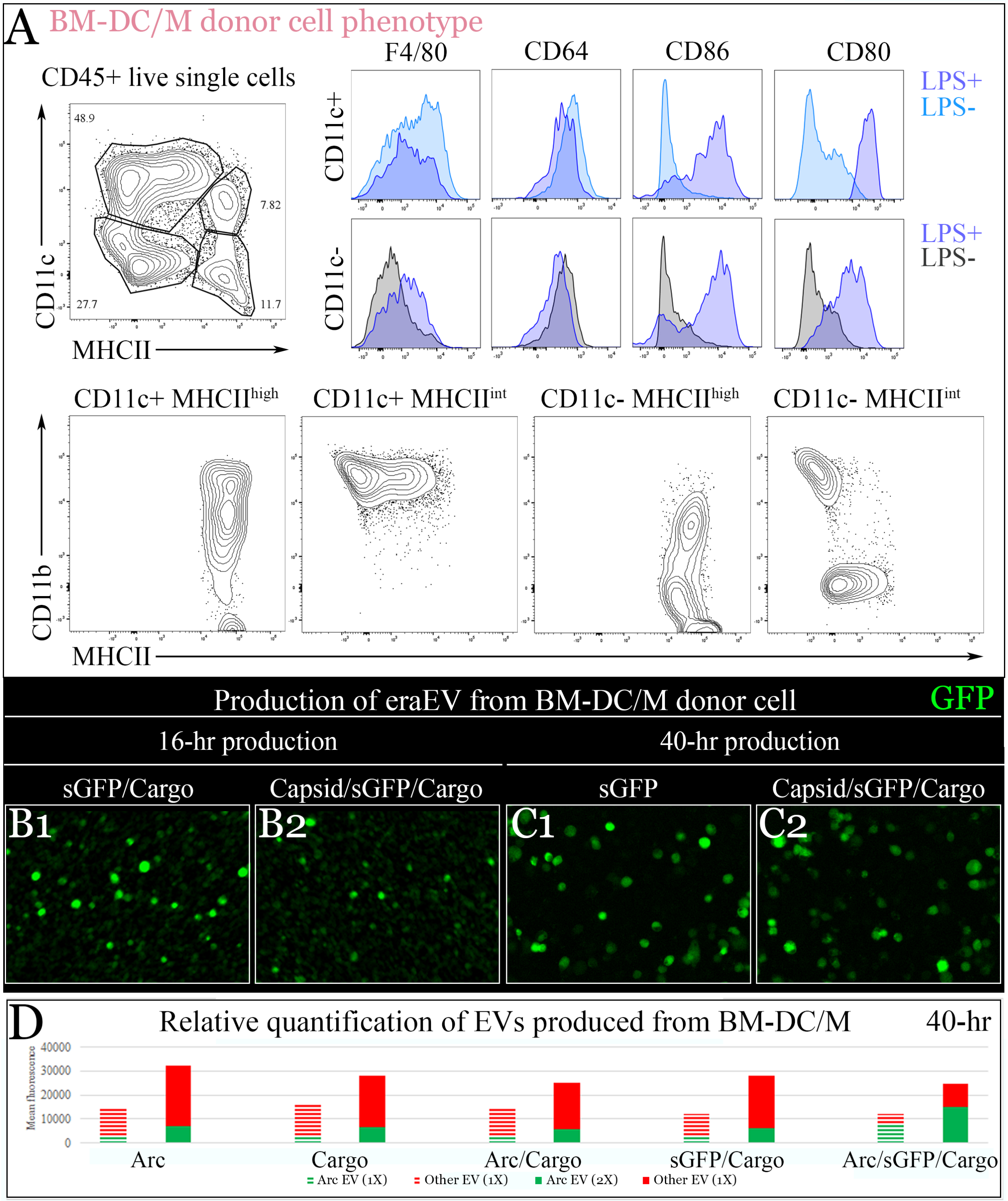


**Fig. S7. Characterization of the self-derived bone marrow differentiated dendritic cell and macrophage (BM-DC/M) as donor cells to produce immunologically inert EVs. (A)** The phenotype of representative GM-CSF/IL4 BM cultures at day 6. CD11c$^{+}$MHCII$^{+}$ BMDCs (C, top left) can be sub-divided based on CD11b and MHCII expression (C, bottom). Boxes depict gates and numbers correspond to percentage of cells in each gate. Histograms showing surface expression of markers by LPS+ MHCII$^{high}$CD11c$^{high}$, LPS+ MHCII$^{high}$-CD11c$^{low}$, LPS- MHCII$^{int}$CD11c$^{high}$ and LPS- MHCII$^{int}$-CD11c$^{low}$ subsets. **(B1-B2)** GFP expression was first lower in the Arc+ group at 16 hours post transfection, but eventually caught up with the Arc– group by 40 hours. **(C1-C2)** To achieve a fine balance between sufficient production of eraEV and abundant encapsulation of cargo, the supernatant culture medium was collected at 40 hours post transfection, when the GFP expression was comparable in donor cells w/ or w/o Arc. **(D)** Sufficient production (> 40 hours) lead to a saturation of EVs in the supernatant culture medium, yielding roughly an equal number of total EVs from each sample group. Meanwhile, the proportion of eraEVs among total EVs is significantly higher in the capsid+/stabilizer+ group. Purified EVs (1X and 2X dilutions) were stained with CMDR and a fluorescent Arc antibody, whose epifluorescence intensity was measured for EV concentrations. Based on this fluorescence reading, we adjusted the concentration of EVs to ensure an equal number of total EVs injected into each animal.

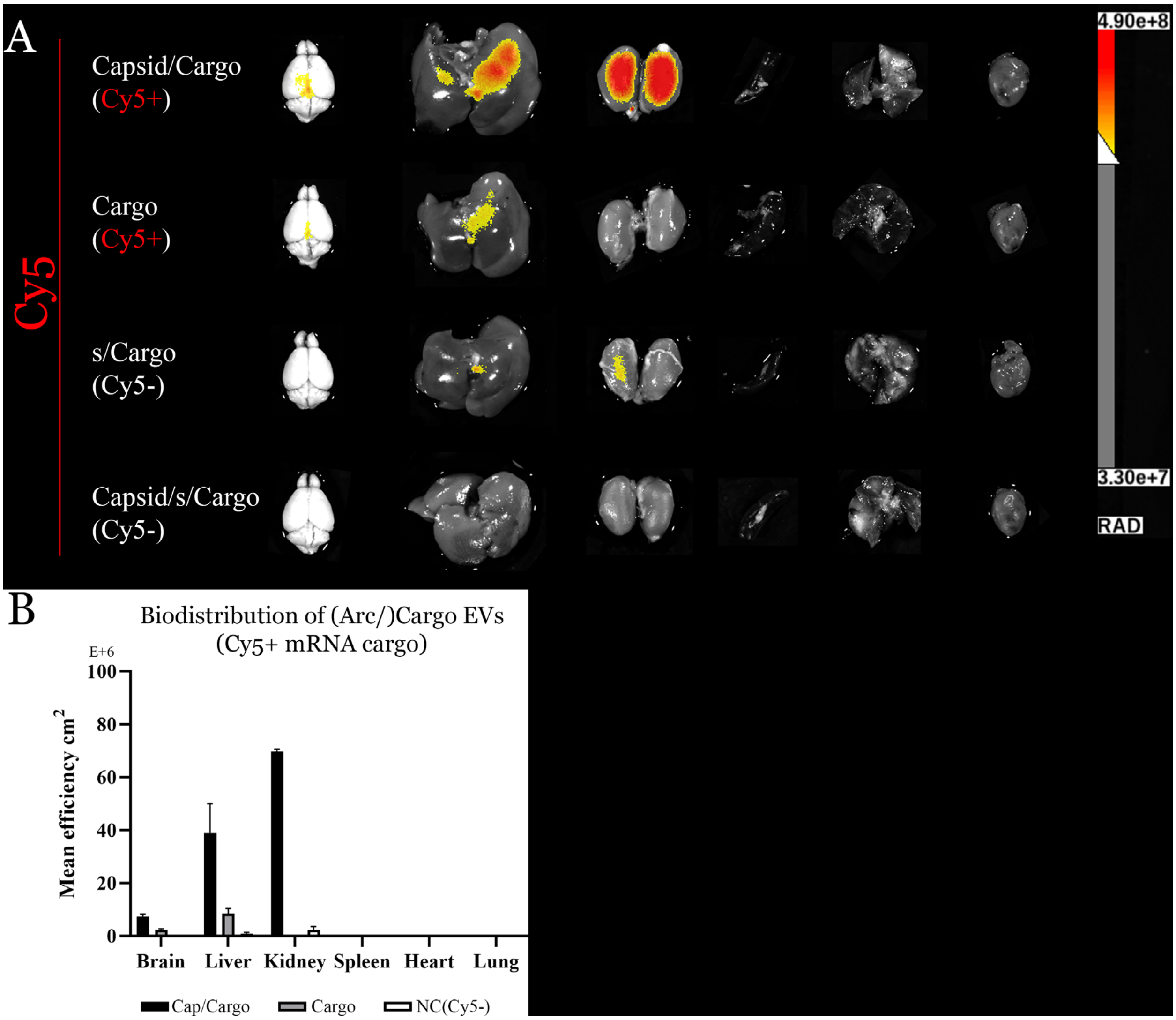


**Fig. S8. Biodistribution of A5U– Cy5+ mRNA delivered by Arc EVs. (A)** IVIS imaging of the *in vivo* biodistribution of Cy5+ cargo mRNA suggest that mRNA cargo does not enrich in the aged brain by Arc EV delivery w/o the A5U stabilizer. A color threshold of 3.5e+8 was applied to subtract the background signal based on the negative control animals (Cy5–/Cy3+ groups, as well as no EV and mock transfection control groups). **(B)** Quantification of the IVIS signal. Mean ± SD, n = 2 in the Arc/GFP group; n=12 from all control groups.

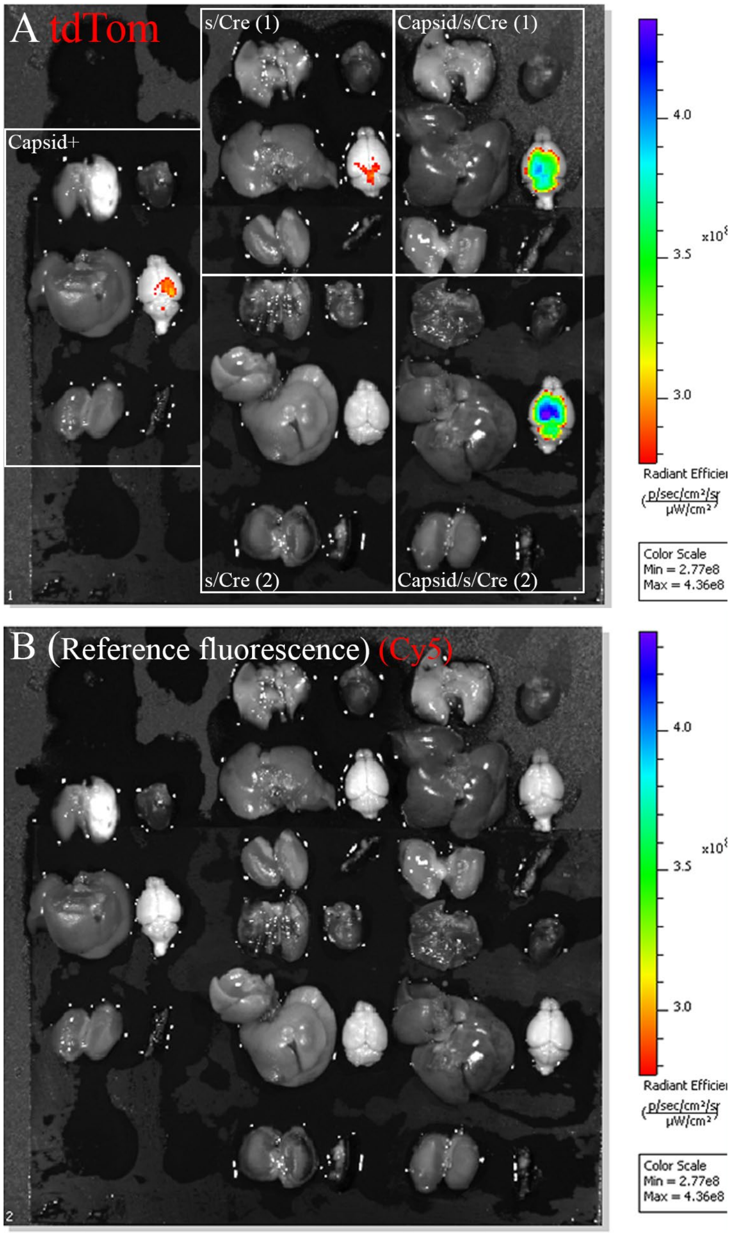


**Fig. S9. Biodistribution of tdTom enabled by Cre+ Arc EVs.** One week after retro-orbital injection, organs were extracted, fixed, and imaged via IVIS, following transcardial perfusion with ice cold DPBS to avoid false positive signal from circulating blood cells. Expression of tdTom at this time point is significantly stronger compared to 3 days after IV injection, as in Fig. 4C1 **(A)**. **(B)** Reference fluorescence was always imaged to ensure specific fluorescent signals.

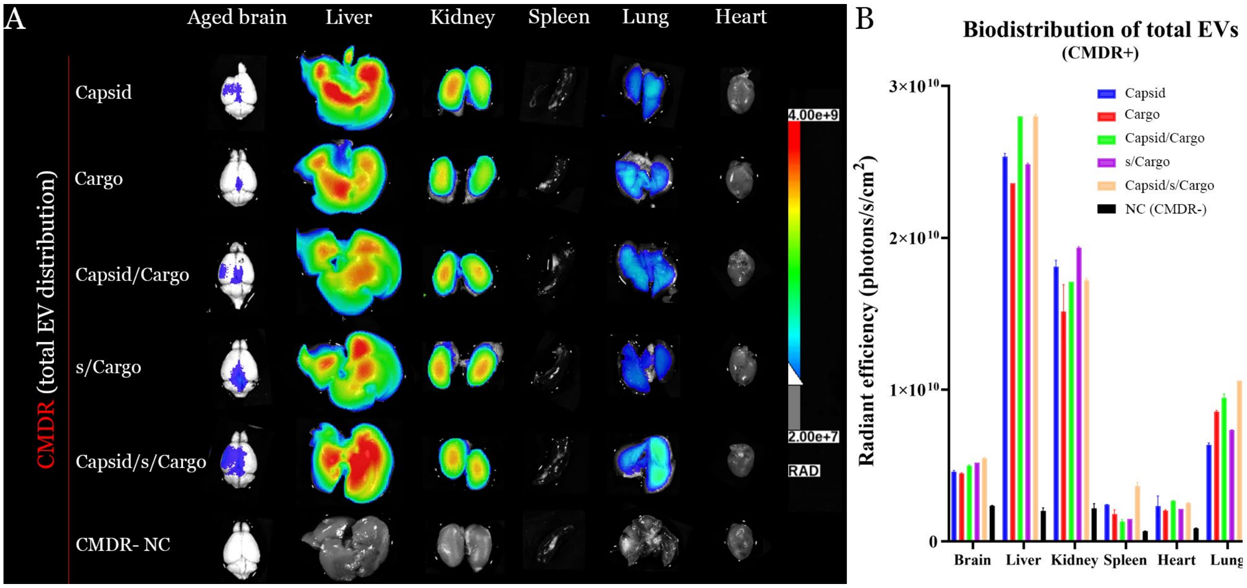


**Fig. S10. Distribution and quantification of total CMDR+ EVs** 3 days after retro-orbital injection of EVs, organs (brain, liver, kidney, spleen, lung, and heart) were extracted from mice following transcardial perfusion by ice cold DPBS, which is to exclude signals from circulating EVs. Organs were then imaged by IVIS. **(A)** Representative images show the biodistribution of total EVs labeled by a plasma membrane dye, CMDR. With the same total number of EVs injected into each control or experimental mouse (per kg body weight), the biodistribution of total EVs was comparable among groups. 3 days post IV injection is the beginning of visible tdTom expression (C1), but by this point CMDR+ total EVs mostly degraded. The biodistribution of CMDR at this time point largely represents the removal of metabolic waste (lipids and CMDR dye). The photo overlay of radiance is displayed, with the color range from 2.0e+7 to 4.0e+9 and a color threshold at 6.0e+8, to subtract the background signal based on the negative control signal level. **(B)** Quantification of the radiant efficiency. Mean ± SD, n = 6 in each group.

**Cy3-sGFP Standard Curve**

| | Total amount of mRNA (ug in 100uL) | Cy3 | Intersection | Slope | R square |
|---|---|---|---|---|---|
| A | 0.1765625 | 62390 | 11463.29 | 178251.4864 | 0.999 |
| B | 0.353125 | 67499 | | | |
| C | 0.70625 | 134414 | | | |
| D | 1.4125 | 263891 | | | |
| E | 2.825 | 494459 | | | |
| F | 5.65 | 1028896 | | | |
| | X | Y | B | A | Y = Ax + B |

| | Mean Fluorescence | Concentration (ug/mL) | Concentration (nM) |
|---|---|---|---|
| NT | 37 | 0.645 | 2.16 |
| Arc | 87 | 0.648 | 2.17 |
| GFP(Cy5)-sGFP(Cy3) | 2128 | 0.762 | 2.55 |
| Arc-GFP(Cy5)-sGFP(Cy3) | 10409 | 1.227 | 4.11 |
| Arc-GFPmodU (Cy3-/Cy5-) | 93 | 0.648 | 2.17 |
| Arc-sGFPmodU (Cy3-/Cy5- | 102 | 0.649 | 2.17 |

MW: 282kDa; Size 877bp

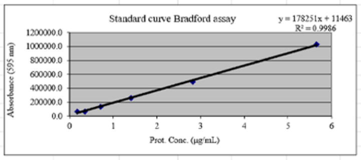


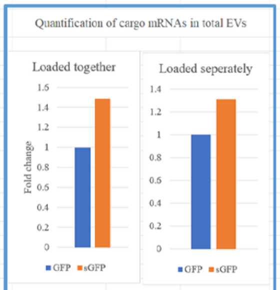


| Loaded together | GFP | sGFP |
|---|---|---|
| Fold change | 1 | 1.483754513 |
| Concentration (nM) | 2.77 | 4.11 |
| **Loaded seperatedly** | GFP | sGFP |
| Fold change | 1 | 1.311345646 |
| Concentration (nM) | 3.79 | 4.97 |

**Cy5-GFP Standard Curve**

| | Total amount of mRNA (ug in 100uL) | Cy3 | Intersection | Slope | R square |
|---|---|---|---|---|---|
| A | 0.15 | 2064 | 152.36 | 12345.4592 | 0.995 |
| B | 0.29 | 2212 | | | |
| C | 0.58 | 6597 | | | |
| D | 1.16 | 17231 | | | |
| E | 2.33 | 28682 | | | |
| F | 4.65 | 57147 | | | |
| | X | Y | B | A | Y = Ax + B |

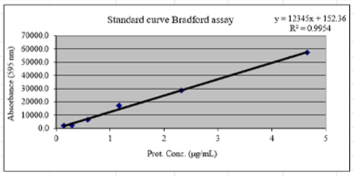


| | Mean Fluorescence | Concentration (ug/mL) | Concentration (nM) |
|---|---|---|---|
| NT | 1 | 0.64 | 0.49 |
| Arc | 1 | 0.64 | 0.49 |
| GFP(Cy5)-sGFP(Cy3) | 129 | 0.65 | 0.94 |
| Arc-GFP(Cy5)-sGFP(Cy3) | 689 | 0.68 | 2.77 |
| Arc-GFPmodU (Cy3-/Cy5-) | 5 | 0.64 | 0.53 |
| Arc-sGFPmodU (Cy3-/Cy5- | 0 | 0.64 | 0.49 |

MW: 232kDa; Size 720bp

**Table S1.** Standard curves and the absolute quantification of fluorescently labelled mRNA cargoes (w/ and w/o A5U) in total EVs. Standard curves were generated by measuring the fluorescence intensity of mRNA transcripts with known concentrations. Both Cy3-sGFP and Cy5-GFP were co-transfected into donor cells to produce EVs, which were then purified with their fluorescence intensity measured. The absolute amount of each cargo in EVs was calculated based on the standard curves. In total EVs with Arc transfected, the ratio between stabilizer+ and stabilizer– mRNA cargoes is 1.48:1.

### Supplementary References


1 Mathieu, M. *et al.* Specificities of exosome versus small ectosome secretion revealed by live intracellular tracking of CD63 and CD9. *Nat Commun* **12**, 4389 (2021). https://doi.org:10.1038/s41467-021-24384-2

2 Lunavat, T. R. *et al.* Small RNA deep sequencing discriminates subsets of extracellular vesicles released by melanoma cells--Evidence of unique microRNA cargos. *RNA Biol* **12**, 810-823 (2015). https://doi.org:10.1080/15476286.2015.1056975
